# The evolutionary history of spines – a Cenozoic arms race with mammals

**DOI:** 10.1101/2023.02.09.527903

**Authors:** Uriel Gélin, Tristan Charles-Dominique, T. Jonathan Davies, Jens-Christian Svenning, William J. Bond, Kyle W. Tomlinson

## Abstract

The role of mammal herbivory in plant evolution is largely unrecognised. Spines on stems are a common and important feature found in ∼9% of eudicot woody plant species worldwide. Spines evolved independently multiple times during the Cenozoic. The timing and extent of spiny plant diversification varied among continents, pointing towards continental rather than global drivers. Spine evolution is closely related to radiation of extant ungulates and extinct ground sloths, rather than climate variation. Diversification began in the Paleogene in herbivore species-rich Eurasia and North America, emerging later in the Neogene in species-poorer South America, Africa and Australia. Spiny lineages expanded their ecological footprint over non-spiny plants, mainly through intercontinental migrations, indicating that spines likely provided a competitive advantage with increasing, and novel, mammal herbivory pressure.

## Main Text

At the end of the 19^th^ century, Charles Darwin referred to the sudden late appearance and rapid diversification of flowering plants in the Cretaceous fossil record (major radiation of eudicots actually ∼160 Mya^1^) as an *abominable mystery* because it opposed his vision of natural selection as a slow and gradual process. Gaston de Saporta drew Darwin’s attention to the fact that this timing coincided with the diversification of insect pollinators, suggesting a process of co-diversification between angiosperms and their pollinators that may have given rise to the huge diversity in flower shape and colour that we see today. Alongside pollinators, insect herbivores also likely played an important role early in the evolution of plants^2^, driving variation in defence-related secondary metabolites^3,4^. Theory on the coevolution of plants and their insect herbivores, and in particular the *escape and radiation* model, provided a powerful framework for the study of plant and arthropod diversification^5^. While the importance of arthropod herbivores in shaping the evolutionary history of plant evolution is now widely recognised, the role of vertebrate herbivores in the radiation of modern plant lineages has received much less attention.

Vertebrate herbivory has been suggested as an important selective force on plant architecture, driving the evolution of structural defence traits^6^–8. Spines are thought to be principally a physical defence against vertebrate herbivores^9^. They are particularly efficient against mammals as they can prick their sensitive mouth parts, and slow feeding rates^10^. Large mammal herbivores can dramatically alter vegetation structure^11,12^, with cascading ecosystem effects^13^, potentially providing an evolutionary pressure that selects for plant defences, as suggested by a close match between the diversification of spiny plants and bovids in Africa around 16 Mya^7^.

Globally, spiny plants are more common in open and arid or seasonally dry ecosystems, such as desert, savanna and shrubland, compared to closed, forest ecosystems^7,14^. The distribution of closed and open ecosystems is typically attributed to abiotic factors such as climate and soil^15^, and this has led to suggestions that the diversification of spiny plants may have instead been precipitated by global drying events associated with global cooling that bound moisture in Antarctic and Arctic ice^16^. However, it is now recognized that large herbivores as disturbance factors strongly shape the balance between open and close vegetation in large parts of the world^17,18^. As the continental expansions of major modern mammal herbivore lineages pre-date major climate drying events, and were asynchronous across regions^19^,it is thus possible to contrast these competing hypotheses by comparing the temporal sequence of events.

Several key climatic events led to global cooling during the Cenozoic, including the opening of the Tasmanian gateway ∼33 Mya^20^ (−4-6 degrees in ∼1my) at the Eocene-Oligocene boundary, and the initial closing of the Central American seaway at ∼15-13 Mya^21^ (−2-6 degrees in ∼2my^22^). Across these same time periods, mammal exchanges between continents have occurred extensively across North America and Eurasia from ∼55 Mya, although they are more recent between Africa-Eurasia (∼20 Mya when the Tethys seaway closed definitively), and between North and South America (∼3.5 Mya^23^). In contrast, Australia has been isolated from the rest of the world since the early Eocene^24^. While continental bridges clearly provided the best opportunity for terrestrial animal migrations, it is likely that they also facilitated the movement of plants^24,25^ even without complete closure^26^. However, plants may also have been better able to disperse long distances (see McGlone *et al*.^27^ for an example of cross-ocean long-distance dispersal) even without land bridges, providing additional opportunities for intercontinental migration.

Here, we explore the worldwide evolution, diversification and biogeography of stem spines in eudicots in relation to global climate change, continental movements and large mammal herbivore. Stem spines include structures modified from the epidermis (prickles), axial meristems (thorns), stipules (stipular spines), and petioles and rachis^28^. We used the large, dated phylogeny of Zanne *et al*.^29^, subsampled for woody species, coded for the presence or absence of stem spines, and employed ancestral state reconstruction to estimate the earliest emergence of stem spines on each continental land mass (excluding Antarctica) and matched this to mammal large-herbivore diversity dynamics. Because both phylogenetic and fossil records are incomplete (owing to the megafauna extinctions during late-Quaternary and earlier, and taphonomic bias^30^–32, respectively), we combined data from both records to reconstruct the diversification of major mammal herbivore lineages, including Afrotheria (origin: Africa), Diprotodontia (Australia), Artiodactyla and Perissodactyla (Laurasia), and Meridiungulata and Pilosa (South America).

Stem spinescence is a common and widely evolved trait, with stem spines present in 23% (*N* = 271) of families and 56% of orders (*N* = 43), cumulatively representing 9% of the 7420 woody eudicot taxa represented in the Zanne *et al*.^29^ mega-phylogeny. North America is richest in plants with stem spines (∼15% of species), followed by Eurasia (∼10%), Africa (∼8%), South America (∼7%) and lastly Australia (∼4%) (table 1, fig. 1). We identified 98 independent evolutionary events of spiny stems occurring all during the Cenozoic, sometimes referred to as “the Age of Mammals”, with the earliest stem spines emerging in the late Paleocene ∼60 Mya (fig. 1).

**Table 1:** Stem spine frequency and date of emergence across continents.

|  | Africa | Eurasia | North<br>America | South<br>America | Australia |
| --- | --- | --- | --- | --- | --- |
| Total number of sampled species on the continent | 1463 | 1864 | 1099 | 1715 | 1279 |
| Number of spiny species | 111 | 208 | 165 | 137 | 49 |
| Proportion of spiny species (%) | 7.6 | 11.2 | 15 | 6.8 | 3.8 |
| **Date of spine emergence (Mya)** | **18** | **61** | **46** | **32** | **18** |

Note: Total number of woody eudicot species sampled in each of five continents (from Zanne *et al.* (29)), the percentage that have spiny stems and estimated dates of emergence of first spiny plant (from ancestral reconstructions).

**Fig. 1:**
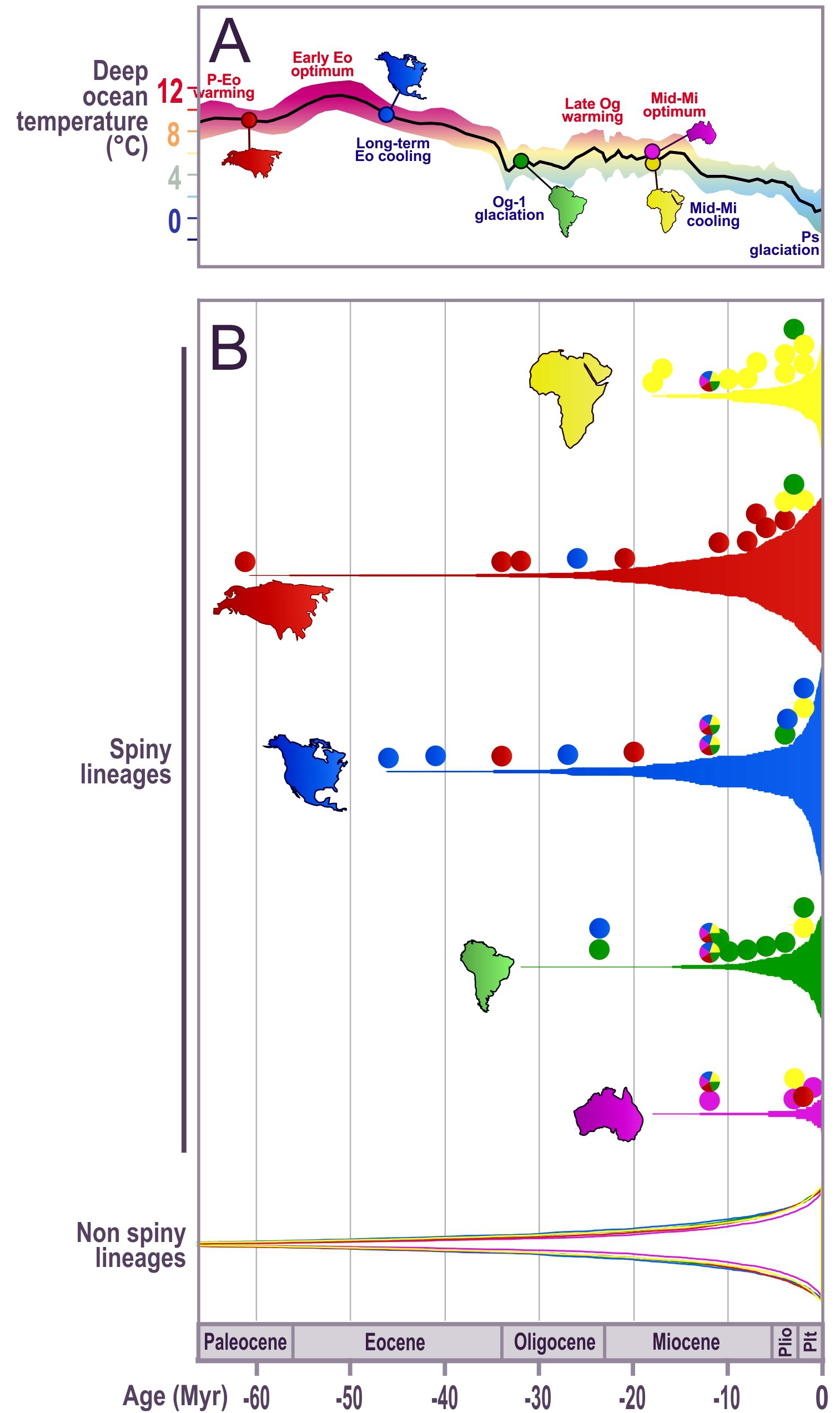
Global evolution of plants with stem spines. Proportion of plant lineages with stem spines within each continent (N=7420; Africa: 7.6%, Eurasia: 10.3%, North America: 15%, South America: 6.8%, Australia: 3.8%) relative to non-spiny plants (bottom, B) and global temperatures. Coloured continent silhouettes in (A) indicate emergence date of stem spines on each continent. Circles above each spiny plant diversification curves indicate independent first emergence of spinescent lineages in a given plant family (either by *in situ* evolution or immigration). Colours of the circles indicate the inferred origin of the spiny clade based on reconstructions using all eudicot woody species (N=9898). The similar timing and dynamic of diversification we observed for non-spiny plant lineages among continents is consistent with the importance of global climate on angiosperm diversification, and contrasts with the complex diversification dynamics of spiny plant lineages.

### (1) Timing of stem spine evolution and radiation was asynchronous

We found large variability in the timing of emergence and radiation of spiny plants among continents, and that the timing of diversification differed between spiny and non-spiny lineages (fig. 1, table 1). The asynchrony in stem spine emergence and timing of diversification peaks strongly suggests that continent-specific drivers, not global factors, selected for spines. While non-spiny angiosperms emerged and diversified during the mid-Mesozoic^33^, spiny plant history is anchored in the Cenozoic (fig. 1B), ∼100 million years later. We estimate that spiny stem species emerged first in Eurasia (Paleocene) and then North America (mid-late Eocene), later in South America (late Oligocene-Miocene), Africa and Australia (early Miocene, table 1, fig. 1 and 2). Spiny fossils have only rarely been reported, but their dates are consistent with our reconstructions pointing to a Laurasian origin of spines: a mid-Eocene (∼50 Mya) specimen from North America^34^, one ∼47 Mya, and then seven morphotypes representing at least five lineages from the late Eocene ∼39 Mya in Eurasia35. However, the number of spiny lineages remained low (comprising <5% of spiny lineages surviving to the present) until major diversification events in the Oligocene in Eurasia, the Miocene in North America, Africa and South America, and the Pliocene in Australia (fig. 1 and 2), ages also characterized by important fauna turn-over. Secondary radiations of spiny lineages followed in the Miocene within Eurasia, and in the Pliocene within North America, Africa and South America (fig. 1 and 2, table S2).

**Fig. 2:**
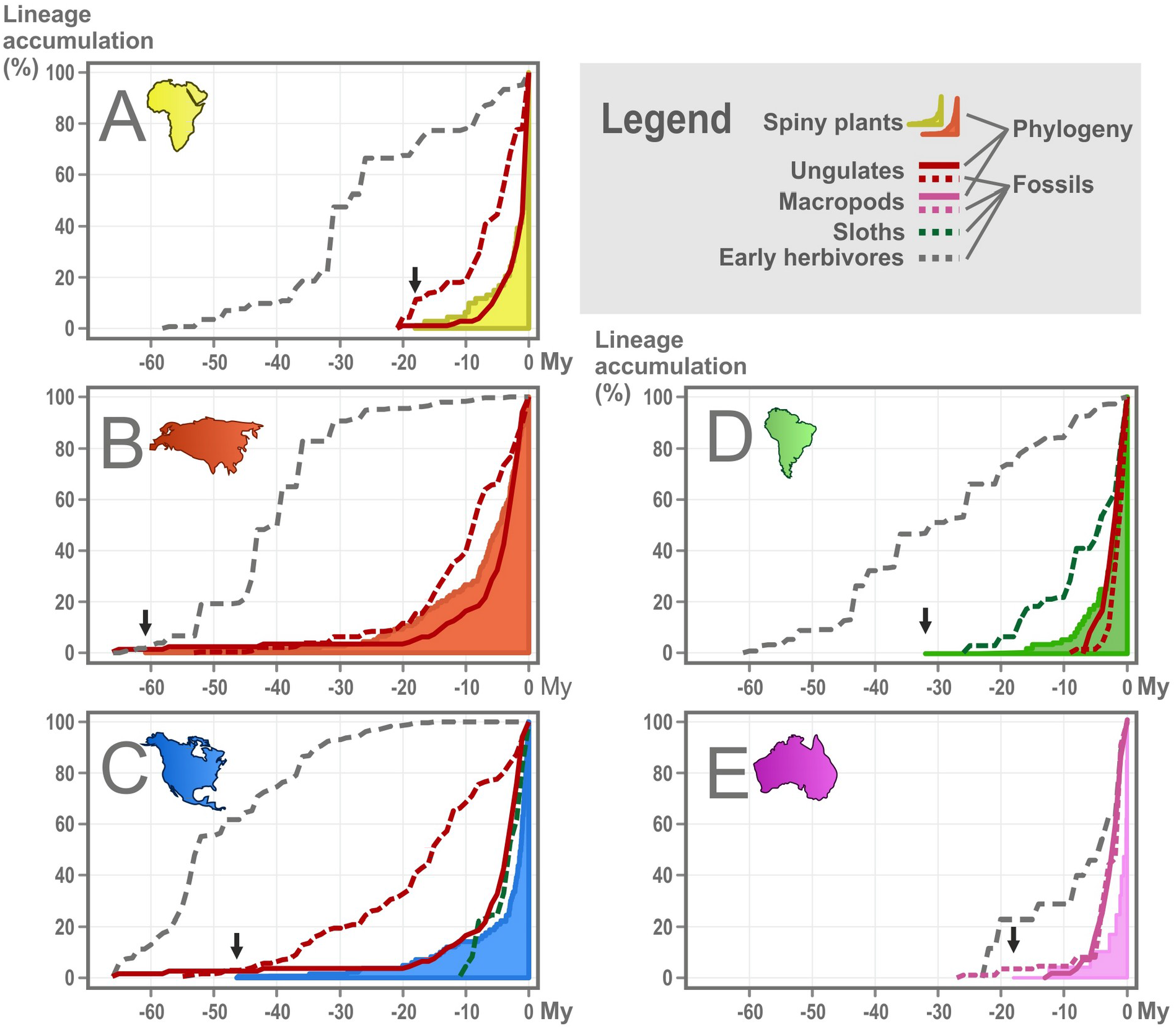
Diversification of large mammalian herbivores and plants with stem spines. Proportion of plant lineages with spiny stems within each continent (N=7420) contrasted with diversity of dominant large mammalian herbivores (N=340 species from phylogeny and N=5645 from fossil records). We show the Eurasian ungulates phylogeny in the plot for North America as important faunal exchanges occurred between these two continents. Black arrows indicate timing of spiny plant emergence in each continent according to phylogenetic reconstruction.

Stem spines emerged and then diversified under both warmer / humid (Eurasia and North America) and cooler / drier (South America, Africa and Australia) global climates (fig. 1A). Although we observe a general negative relationship between angiosperm diversification and global temperature, the strength of this relationship was notably weaker for spiny plants in Africa, Eurasia and South America relative to that observed for non-spiny plants (tables S3-4), suggesting that global cooling was not a major driving force selecting for spines in these continents. However, spiny plant diversification was more sensitive to global temperatures in both the relatively isolated Australian continent, and herbivore-rich North America (tables S3-4). Spines may be adaptations to arid environments in some clades (e.g., Cactaceae^36^); however, temperature likely also impacts herbivore movements, more obviously through its effect on sea level and the emergence of intercontinental land bridges such as Beringia which facilitated fauna and flora exchanges between Eurasia and North America^37^.

### (2) Stem spine evolution and radiation is correlated with the diversification of mammal lineages

We found that the diversification of spiny stem plant lineages, reflected by their current continental abundance (table 1), matches closely to the diversification of mammal herbivores across the globe. The fit of the model explaining the variability in diversification rates for Africa, Eurasia and South America including the extant Laurasian ungulate families (Artiodactyls and Perissodactyls) is vastly preferred over the model with global climate (fig. 2; tables 2 and S3-4). While climate was also an important predictor of spine evolution in Australia and North America, diversity of large mammal herbivores was still positively correlated with spiny plant diversification. Ungulates belonging to extant families in North America as well as some now extinct species in distinct orders, such as sloths in North and South America, and Afrotherians in Asia, appear to have triggered the evolution of stem spines (tables S3-5). However, we find little or no evidence of spiny stem co-diversification with Afrotherians in Africa, *early* Laurasian ungulates in North America and Eurasia, meridiungulates (in North America and in South America, nor with either macropods or non-macropod Diprotodonts in Australia (tables S4).

**Table 2:**
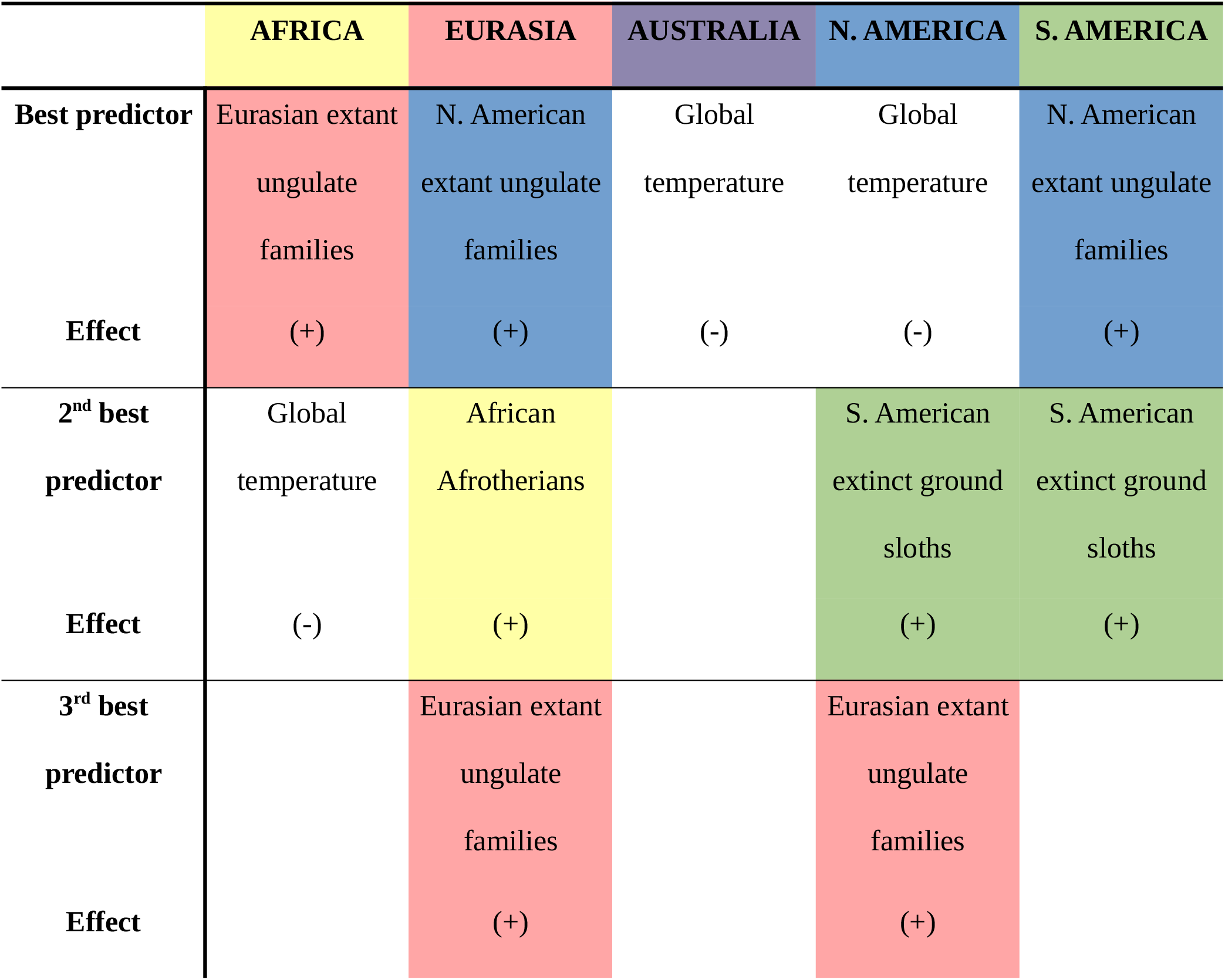
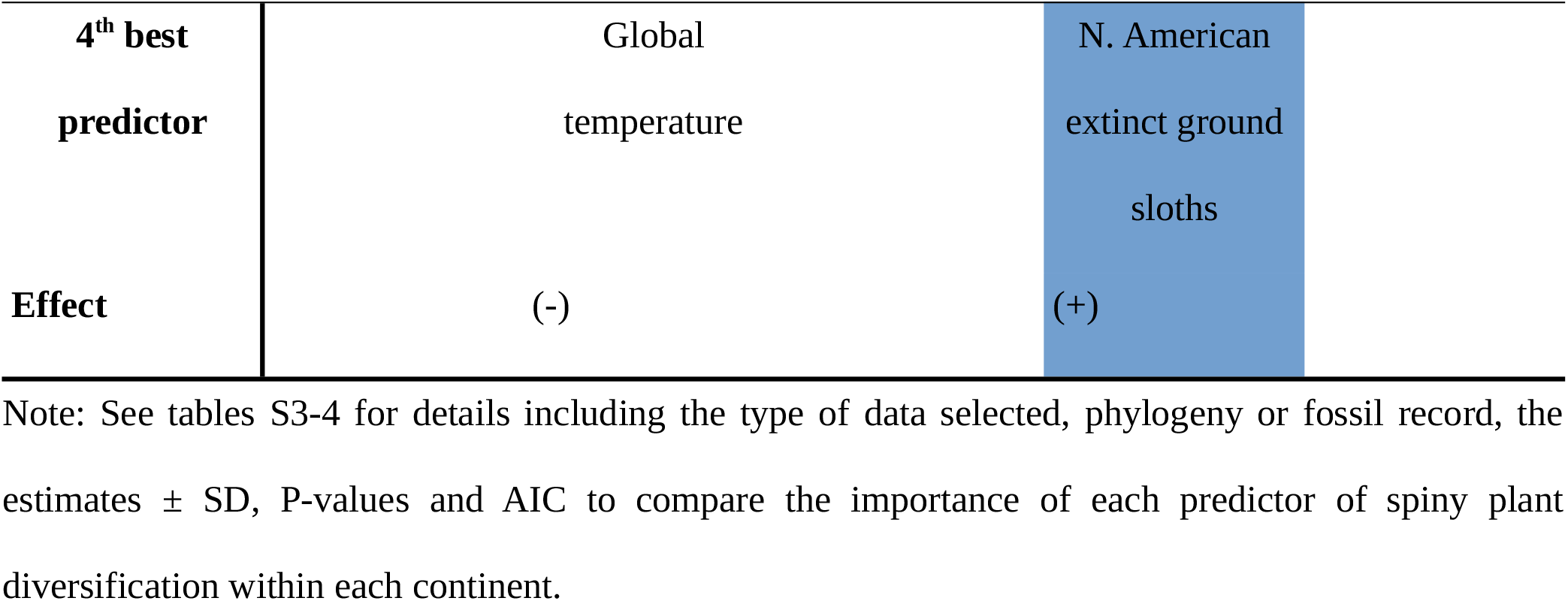
Summary of the main significant predictors of spiny plant diversification within each continent, ranked from most to least important. Spiny plant diversification generally decreased with temperature, possibly suggesting that herbivores imposed a greater selective pressure favouring spines during periods of low temperature. However, it was more sensitive to global temperatures in both the relatively isolated Australian continent, and herbivore-rich North America, probably linked to the importance of plant and/or herbivore migration facilitated by low sea level when water was locked up in polar icecaps. Coloured background indicates the importance of inter-continental mammalian herbivore migration affecting spiny plant radiation in each continent (yellow for Africa, red for Eurasia, purple for Australia, blue for North America and green for South America).

South America, Africa, and Australia harbour more recent and modest spiny stem plant and mammal herbivore diversity compared with North America and Eurasia that were exposed to substantial herbivory pressure from large herbivore assemblages with very diverse ways of feeding (e.g., camelids, ruminants, horses, rhinos, ground sloths) during most of the Cenozoic (table S5 and fig. S1). Eudicot spiny plant radiation is strongly associated with the diversification of large mammal herbivores, especially modern ungulate families, with the exception of Australia, the only continent without native ungulates, confirming globally the trend previously identified for Africa^7^. However, whereas, within these Laurasian ungulates, bovids likely selected for the evolution of stem spines in Africa, camelids, equids, cervids and other non-bovids with similar diets (ranging from pure browsers to pure grazers) to those of the modern-day African savannah ungulates^38^, were likely more important selective pressures in South and North America, notably because bovids never reached South America and were late-arriving, predominantly grazers in North America^38,39^. Yet, as the other clades, early Laurasian ungulates do not appear to have imposed a strong pressure favouring the evolution of stem spines, perhaps due to their size, lack of feeding specialization and low population densities. Australia is well represented by Macropod herbivores, but has less than half the proportion of spiny stem plants recorded in Africa, further suggesting that spiny stems and possibly physical defences may have been, as for other Diprotodont families, a poor deterrent to kangaroo-like herbivore that have very different way of feeding than ungulates.

We found a stronger impact of migrant herbivores, such as North American ungulates in Eurasia, South American sloths in North America and Eurasian ungulates in Africa, relative to local fauna (tables 2 and S3-4), perhaps reflecting the importance of novel herbivore pressure in driving the diversification of spiny lineages and offering some explanations for the possible effect of Afrotherians in Eurasia but not in their continent of origin. We suggest, therefore, that spiny plants and mammal browsers may have engaged in a co-evolutionary arms race that played out asynchronously across continents, and resulted in the co-diversification of spiny plant lineages and ungulate herbivores, mediated by regional climates in some continents.

#### Climatic constraints and herbivory

Vegetation structure in dry climates is particularly sensitive to large-herbivore effects (18) while drier and more open environments also tend to support greater diversity and abundance of spiny species and herbivores in the present day^7,40^. We found that spiny plant diversification generally decreased with temperature and a significant interaction between herbivory pressure and temperature suggests that herbivores imposed a greater selective pressure favouring spines during periods of low temperature (tables S3-4). This interaction might help explain why spiny lineages were apparently favoured during cooler climates in North America, which was rich in large ungulate herbivores, and in Australia, host to a unique mammal herbivore community. Lower sea level occurring when water was locked up in polar icecaps during periods of low temperature, notably linked to the presence of similar biomes with convergent climate (dry, seasonally dry) and herbivore (mammal browser) regimes, may have additionally facilitated the exchange of fauna and flora lineages between continents^24^, leading to strongly temperature-related spiny plant radiation.

### (3) Spiny lineages are over-represented in intercontinental exchanges

We show that spiny stem plants are disproportionately represented in inter-continental floristic exchanges (fig. 3), suggesting that mammal browsers likely posed an obstacle to non-native plant establishment, which favoured colonization by plants with spines. Migration of spiny plants between continents has been more common than the *in-situ* evolution of stem spines, and we estimate that between 72% to 97% of extant spiny plants within a continent descend from spiny ancestors that originated on a different continent. The frequency of migration events expanded over time, likely a result of land bridges between formerly disconnected continents (table 1, fig. 1B). Surprisingly, however, the continents where stem spinescence emerged first (Eurasia and North America) were not the main source of spiny clades toward other continents (contributing just 9% and 21% of spiny plants, respectively) but were the main receivers (38% and 31% of spiny plants, respectively for Eurasia and North America) (fig. 3). Australia and South America, which were isolated for most of the Cenozoic, had the highest proportion of plants with stem spines resulting from *in situ* evolution (31% and 25%, respectively), while Africa and South America were the main donor continents. The rich spiny plant flora of South America, in addition to its richness of large herbivores with diverse ways of feeding (e.g., ungulates, sloths, mastodont-like Meridiungulata; table S5 and fig. S1), no doubt explains its importance as a global source of spiny plant lineages. However, Africa has a lower richness of plants with stem spines to Eurasia, yet the African spiny flora appears more adept at colonising elsewhere.

**Fig. 3:**
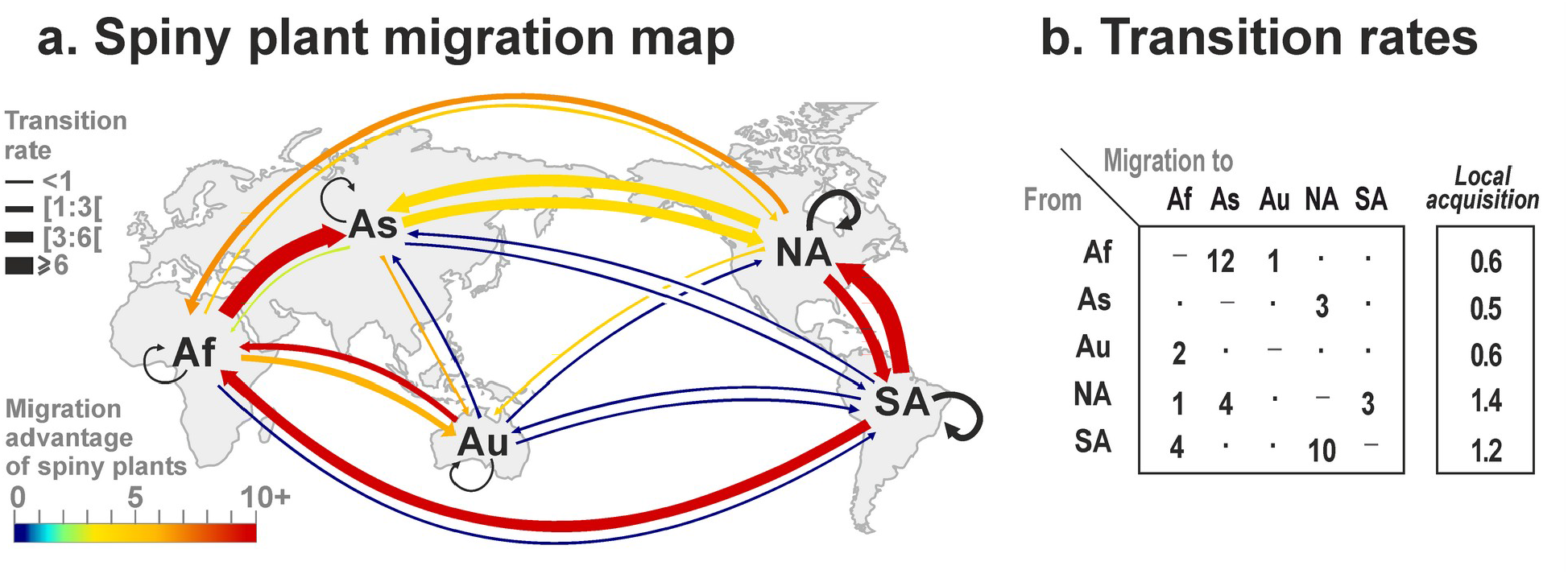
Biogeographic reconstruction of dispersal and evolution of plants with stem spines. (a). Colours of arrows connecting continents represent the relative migration rate for spiny plants relative to total woody plant migration between the same continents. Arrow widths represent the relative global importance of each migration pathway. Loop within continents (black arrows) represents local changes in state from non-spiny to spiny. (b) Transition rate values (X1000). Geographic areas considered for biogeographic reconstructions are: Africa, Eurasia, Australia, North America and South America.

### A substantial but underestimated impact of mammal herbivores on plant evolution

Our classification of spiny stems does not describe the full complexity of structural defence strategies. The leaves can themselves be spiny and it is clear that woody plants have evolved complex branching architectures (“cages”^41^) that also reduce feeding rates by mammals^8^. In addition, our pooling of herbivore types does not capture the nuances of dietary specialization and ways of feeding. Alternative defence types such as chemical ones, also influence mammal feeding behaviour^42^ but, unlike stem spines, are not easily separated from defence against insect herbivores^43^. Nonetheless, we are able to demonstrate the selective effect of mammal browsing pressure on plant trait evolution and radiation.

We provide evidence that extant families of large mammal herbivores, including Laurasian ungulates and sloths, were likely the main selective force driving the global evolution and diversification of spiny plants. Now extinct mammal herbivore lineages and global climate events appear to explain less of the present-day diversity of spiny plants. The local acquisition of stem spines contributes strongly to spiny plant diversity on continents that were relatively isolated from floras elsewhere, but the vast majority of spiny stem plants descend from ancestors that evolved stem spines on other continents. The early timing of spiny plant diversification (table 1), geographically biased transition and migration rates between continents (fig. 3, tables S1-2), as well as ancient mammalian herbivore history (fig. 2, tables S3-5 and fig. S1), all point towards Eurasia as a primary source of spiny plant emergence and diversification.

We suggest an evolutionary arms race between plants and mammalian herbivores played out across the globe in the Cenozoic, favouring the occurrence of stem spines that now adorn at least 9% of woody plants, shaping ecosystem evolution, and plant geographical patterns. The effect of mammals on plant architectural and chemical evolution is likely much more multifaceted, extending to an even greater number of plant lineages^44^.

## Methods

We focus here on trends in net diversification and speciation rates; extinction rates are hard to estimate reliably from phylogenetic trees^45^, and plant evolutionary history, contrary to animal history, does not show strong evidence of mass extinction among higher taxa, but rather alternates between plateaus and bursts of speciation^25^, suggesting that extinction is not the main driver of plant macroevolution^46^. We also expected innovation (speciation rather than extinction) to be both the most important driver and response linking herbivores and spiny plant diversification. Coevolution is then expected to shape the adaptive peaks of interacting clades^47^, here between large mammalian herbivores and spiny plants. We used (Number of spiny lineages at time t+1)/(Number of spiny lineages at time t) for every million year intervals for each continent as a proxy for diversification rate for fossil / phylogenetic data.

### a. Evolution and diversification of stem spines

We reconstructed the evolutionary history of stem spines across 7420 species of woody eudicots represented on the mega-phylogeny of plants from Zanne et al^29^ using the make.simmap functions in phytools^48^ and the ape^49^ and Geiger^50^ R-libraries, assigning each species to one of five geographic continents. We reconstructed the ancestral states for continent of origin and excluded species from subsequent analyses when their region of origin was ambiguous. To control for the relatively recent origin of stem spines and diversification of spiny plant lineages, we allowed rates to vary through time (parameter Δ; Tables S1-2), and constrained evolutionary transitions to preclude reversals from spiny to non-spiny states, reflecting the rarity of this evolutionary event, and avoiding potential bias due to the over-representation of non-spiny lineages among woody taxa^51^.

### b. Evolution of large mammal herbivores

We explored the coincidence between the diversification of mammal herbivores, as estimated from the fossils record or phylogeny of extant species, and the diversification of spiny plant lineages.

#### Phylogeny

We used the mammal phylogeny of Meredith et al^52^ to estimate timing and diversification rate of Artiodactyls-Perissodactyls in Africa, Eurasia, North America, South America (respectively, N=89, N=96, N=33, N=65) and Macropodidae in Australia (N=57).

#### Fossil record

We constructed a dataset from the fossil record^53^ of first occurrence of mammal herbivore species, using the mean date estimate to avoid bias due to extreme values. Our dataset was informed by fossils of hoofed mammals (and ground sloths in the Americas) from Africa (N=438), Eurasia (N=1560), North America (N=2704), and South America (N=822), and Diprotodontidae in Oceania (N=121). For Artiodactyla and Perissodactyla in Eurasia and North America, we distinguished between ancient and migrant or modern lineages, the latter includes all extant native families.

### c. Testing relationships between the diversification of spiny plants, mammal herbivores and global temperatures

We used linear models to determine whether diversification rates of spiny and non-spiny plants at continental level, estimated on the plant mega-phylogeny of Zanne et al29, were rather associated with the diversification rates of mammals estimated from the phylogeny of Meredith et al52 and from the fossil record53, or with global temperature of past environments54. We truncated the numbers to integer values when pooling diversification rates per million years for analyses. To investigate the importance of intercontinental fauna exchanges on plant diversification, we tested for the potential effects of both local and neighbouring continent herbivores. All variables were log transformed to meet model assumptions. First, we ran simple linear models for each predictor of each plant type (spiny, non-spiny) per continent, and then compared the models by AIC55 (Table S3). Second, we ran full multilinear models with all predictors for each plant type (spiny stem, non-spiny stem) per continent, to test whether different predictors explained complementary components of species diversification. Third, we tested interactions between predictors, sub-setting the data to only include data starting from the first occurrence of the herbivore group being tested, to test if global temperature impacted the effect of the herbivore group on spiny plant diversification. Sample size thus varied among the interactions tested and was equal to age of first occurrence+1 (Tables 4). North American sloth and South American ungulate diversification were highly correlated (r = 0.98) so they were tested separately in the model for South American spiny plant radiation. All analyses were performed in the R v4.1.2 statistical programming environment (R Foundation for Statistical Computing, Vienna, Austria)

### d. On the reliability of phylogenies and the fossil record

Both fossil and phylogenetic rate estimates are susceptible to biases. Geographic bias in the taphonomic record can skew fossil rate estimates, as illustrated, for example, by the absence of early Diprotodontidae in the Australian fossil record before the Oligocene. Phylogenetic estimates can be sensitive to the particular tree topology used^29,56,57^. However, the dates of plant orders in the Zanne et al. phylogeny are broadly congruent with estimates from other sources (Angiosperm Phylogeny Website, www.mobot.org), and alternative phylogenetic reconstructions do not suggest age estimates are systematically biased either too old or too young^7^. Critically, when data on both mammal phylogeny and fossils where available, they generally indicated similar pace and timing of herbivore diversification (Africa, Eurasia and Australia), although fossils of ungulates in Africa and Eurasia and especially North America indicated slightly higher diversification rates than those estimated from their phylogenies.

## Aknowledgements

Thanks to the main contributors to the data we used from the Fossilworks database: C. Clyde, G. J. Prideaux, L. M. Abraczinskas, A. D. Redline, K. D. Rose, E. I. Belyaeva, C. B. Schultz and C. H. Falkenbach, J. M. Harris, F.H. Brown, and M.G. Leakey, R. Secord, A. J. Kihm, M. D. Leakey and J. M. Harris, J. de Heinzelin., J. Sudre, H. B. Wesselman, M. G. Leakey, D. M. Schankler, J. A. Holman, D. C. Fisher, and R. O. Kapp, M. Bon, G. Piccoli, and B. Sala, C. S. Churcher and P.E.L. Smith, G. D. van den Bergh. Thanks to Catherine Menvielle for helping with spine coding.

## Funding

National Natural Science Foundation of China grant 31470449 (KT)

Yunnan Province Thousand Talents Plan grant number E0YN021 (KT)

Chinese Academy of Sciences and The World Academy of Sciences (CAS-TWAS)

President’s Fellowship for International postdoctoral students grant 2016PB011 (UG)

China Postodoctoral Science Fund grant 2016M600755 (UG)

VILLUM Investigator project “Biodiversity Dynamics in a Changing World” funded by VILLUM FONDEN grant 16549 (JCS)

Independent Research Fund Denmark | Natural Sciences project MegaComplexity grant 0135-00225B (JCS)

## Author contributions

UG, KT, TCD and JD designed the project.

UG developed the idea and collected the data.

UG designed the model and analysed the data with the support from TCD and JD

UG wrote the manuscript in consultation with KT, TCD, JD, WB and JCS

UG, KT, TCD, JD, WB and JCS provided critical feedback and helped shape the analysis and final manuscript.

## Data and materials availability

The datasets generated during and analysed during the current study are available from UG on reasonable request.

## Supplementary Materials

Figs. S1 to S2

Tables S1 to S5

**Fig. S1:**
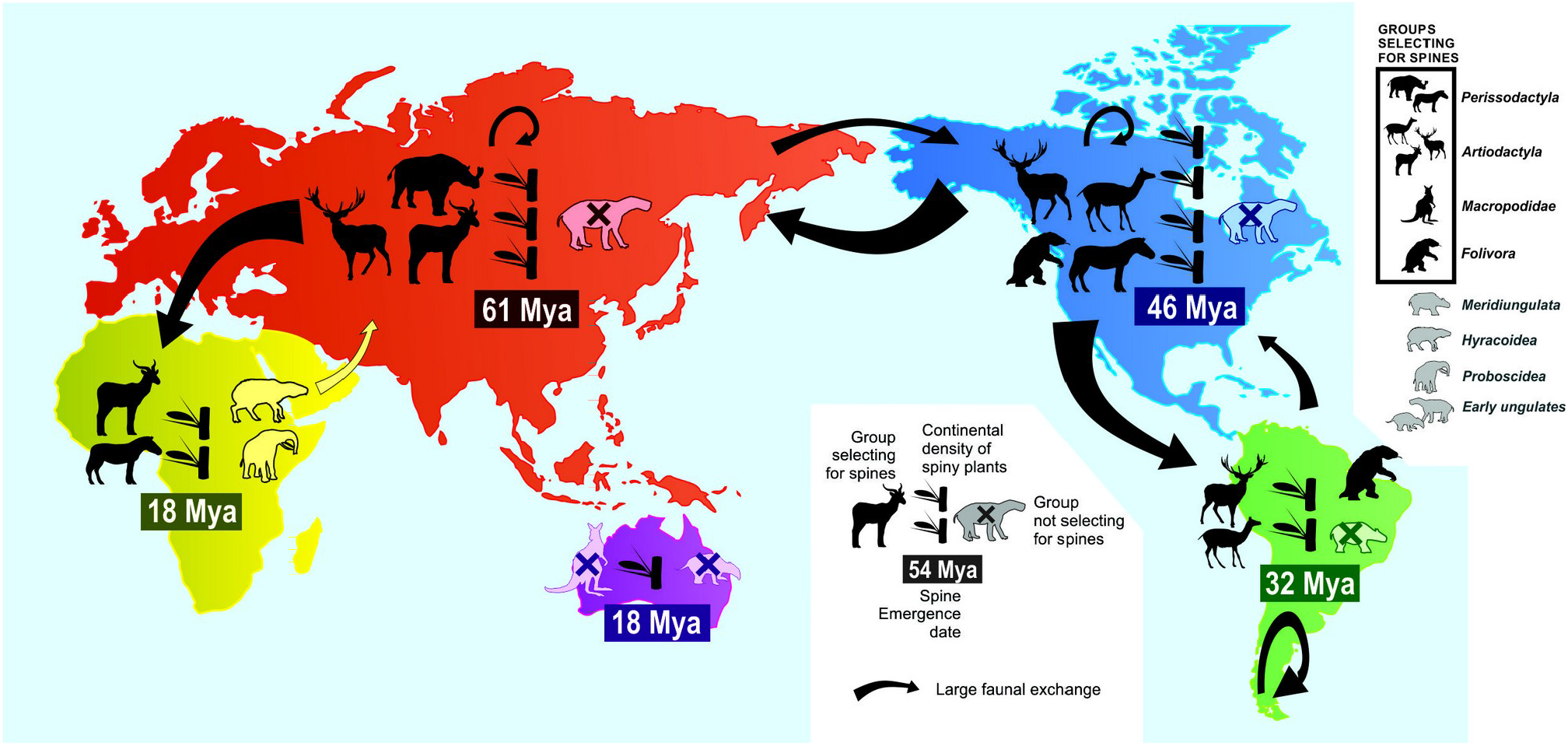
Continental evolution and abundance of spiny plants has been mainly driven by speciation, migration and diversity of mammalian herbivores, especially Artiodactyla/Perissodactyla, and in North America, Folivora.

**Table S1:**
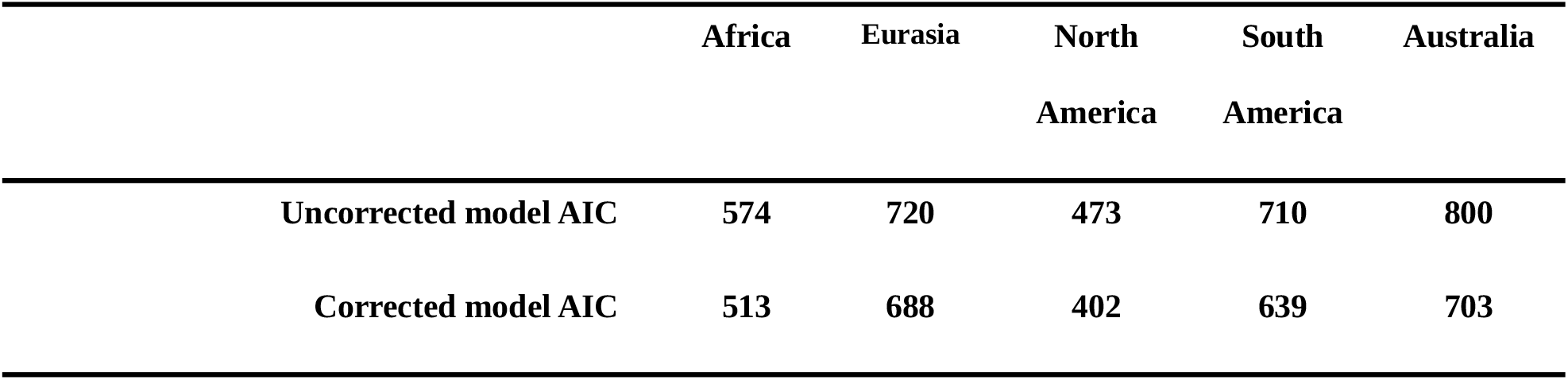
Importance of correction for inflation of diversification rate (parameter Δ) in each continent. Corrected models including Δ always fitted the data better.

**Table S2:** Relative fit of models fitting diversification rates to each continent (rows) using the correction for diversification inflation, Δ, either from the same continent or each of the other continents (columns); AIC values are given with AIC weights in parentheses. Emboldened values indicate best models in each row. Africa and North America show similar inflation of diversification rates, but other continents are best explained by their own Δ values.

|  | **Africa ($\Delta=17$)** | **Eurasia ($\Delta=9$)** | **Australia ($\Delta=60$)** | **North America ($\Delta=15$)** | **South America ($\Delta=26$)** |
| --- | --- | --- | --- | --- | --- |
| **Africa** | **513** (0.45) | 520 (0.02) | 538 (0.00) | **514** (0.41) | 516 (0.13) |
| **Eurasia** | 699 (0.00) | **688** (0.97) | 790 (0.00) | 695 (0.03) | 718 (0.00) |
| **Australia** | 416 (0.00) | 430 (0.00) | **402** (0.98) | 418 (0.00) | 411 (0.02) |
| **North America** | **639** (0.43) | 644 (0.04) | 667 (0.00) | **639** (0.48) | 643 (0.06) |
| **South America** | 706 (0.18) | 722 (0.00) | 716 (0.00) | 708 (0.07) | **724** (0.75) |

**Table S3:**
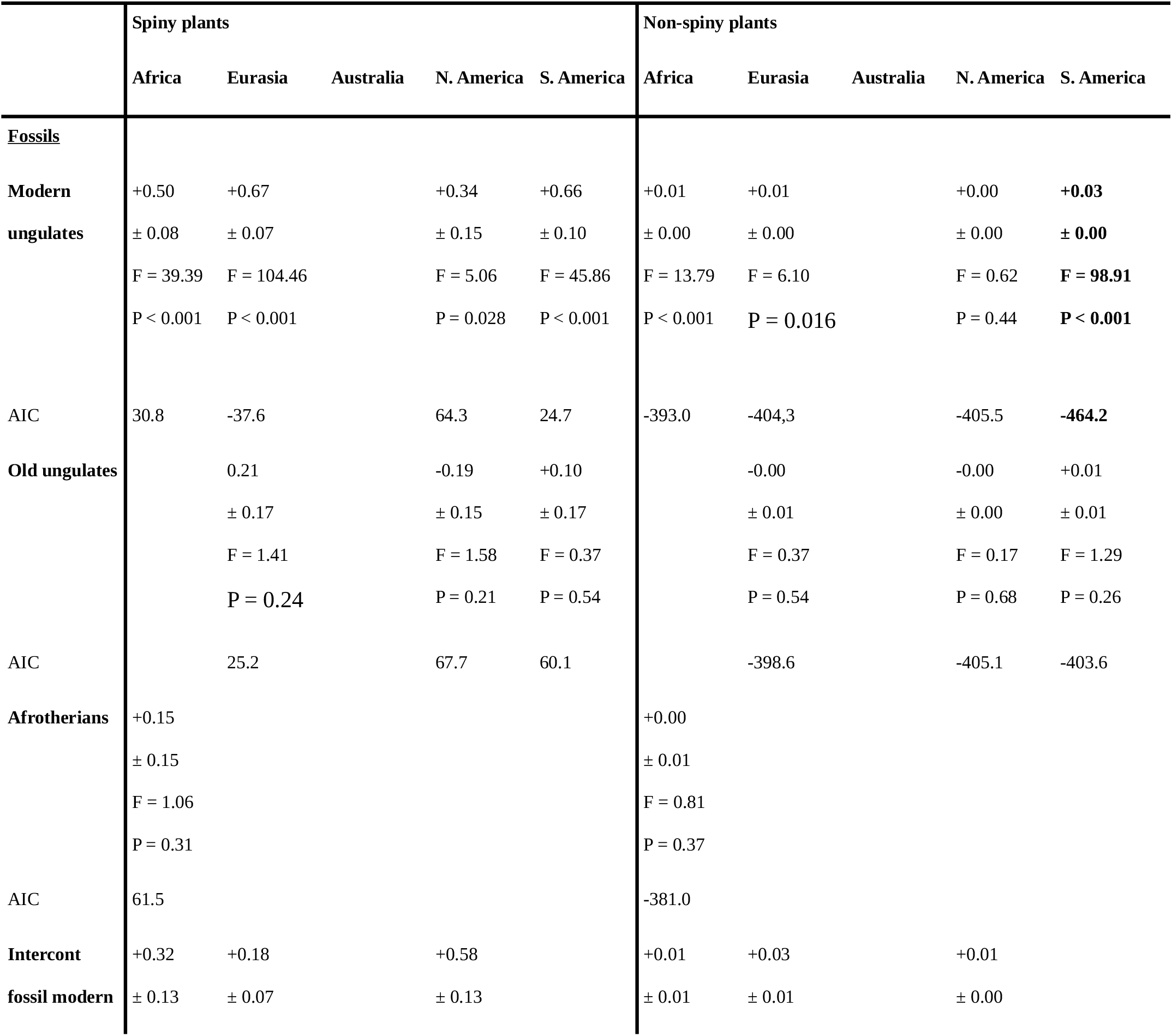
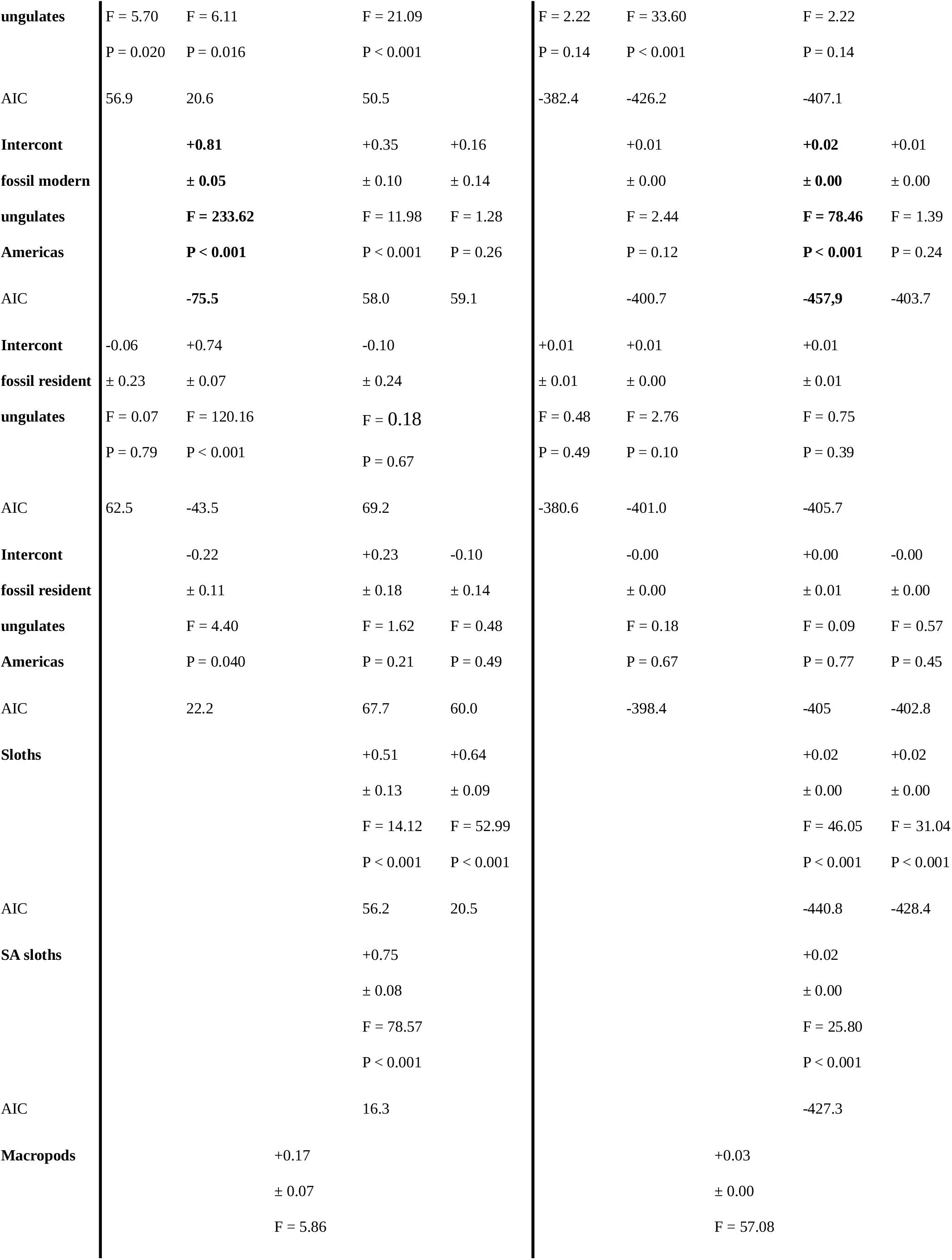
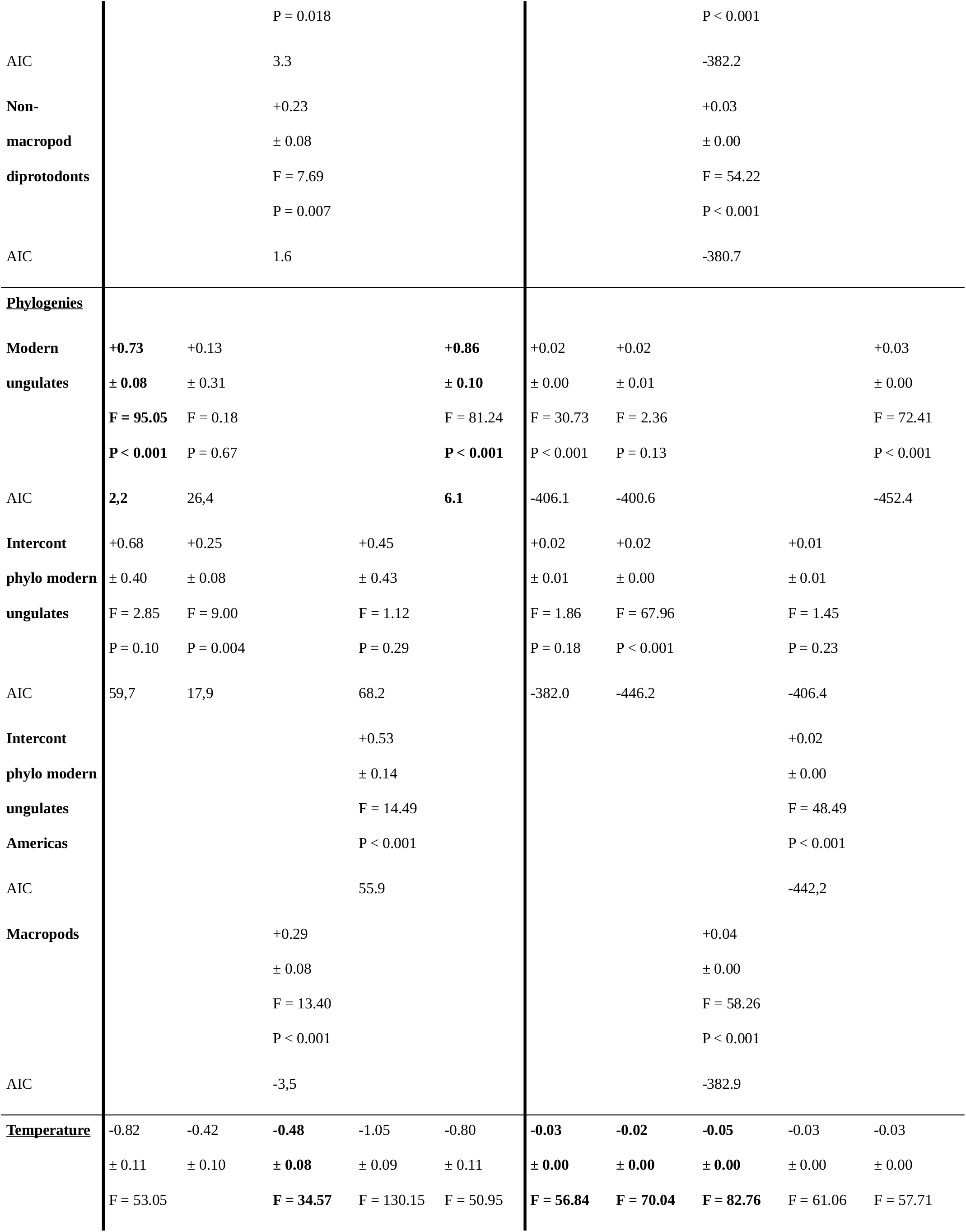
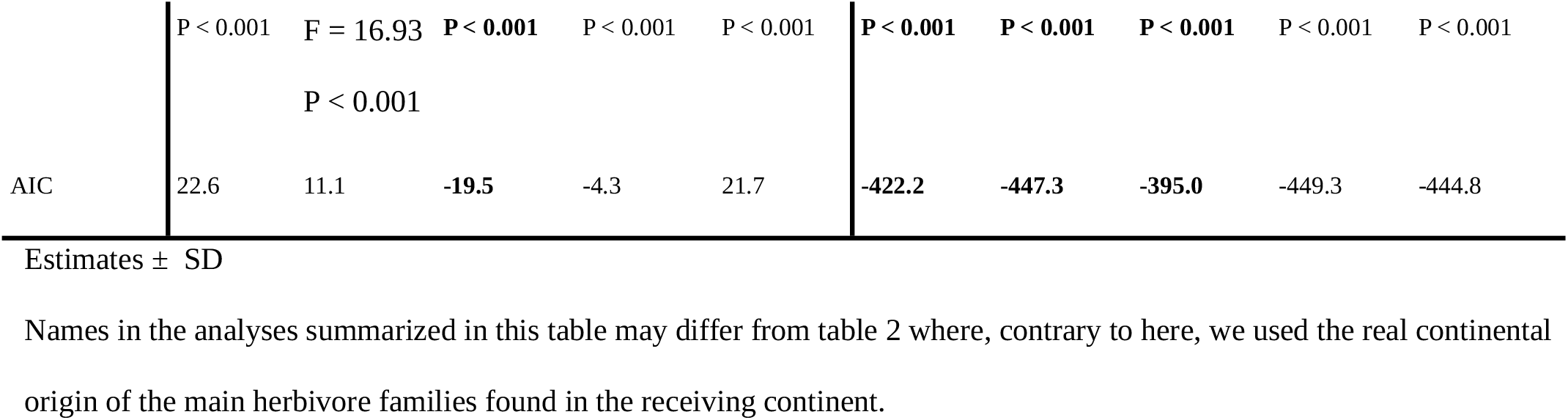
Estimates of linear models and AIC values testing the relationship between herbivore and plant (spiny and not spiny) yearly diversification rate and global temperature for the Cenozoic in 5 continents, one variable at a time (N = 67). Bold indicates the model fitting the best the data. Diversification of herbivores generally fit better to spiny plants while temperature fits better to non-spiny plants but the opposite pattern was found in Australia and North America; “migrant” herbivores appearing the more closely linked to spiny plant diversification but South American sloths and African Afrotherian may have also been involved.

**Table S4:**
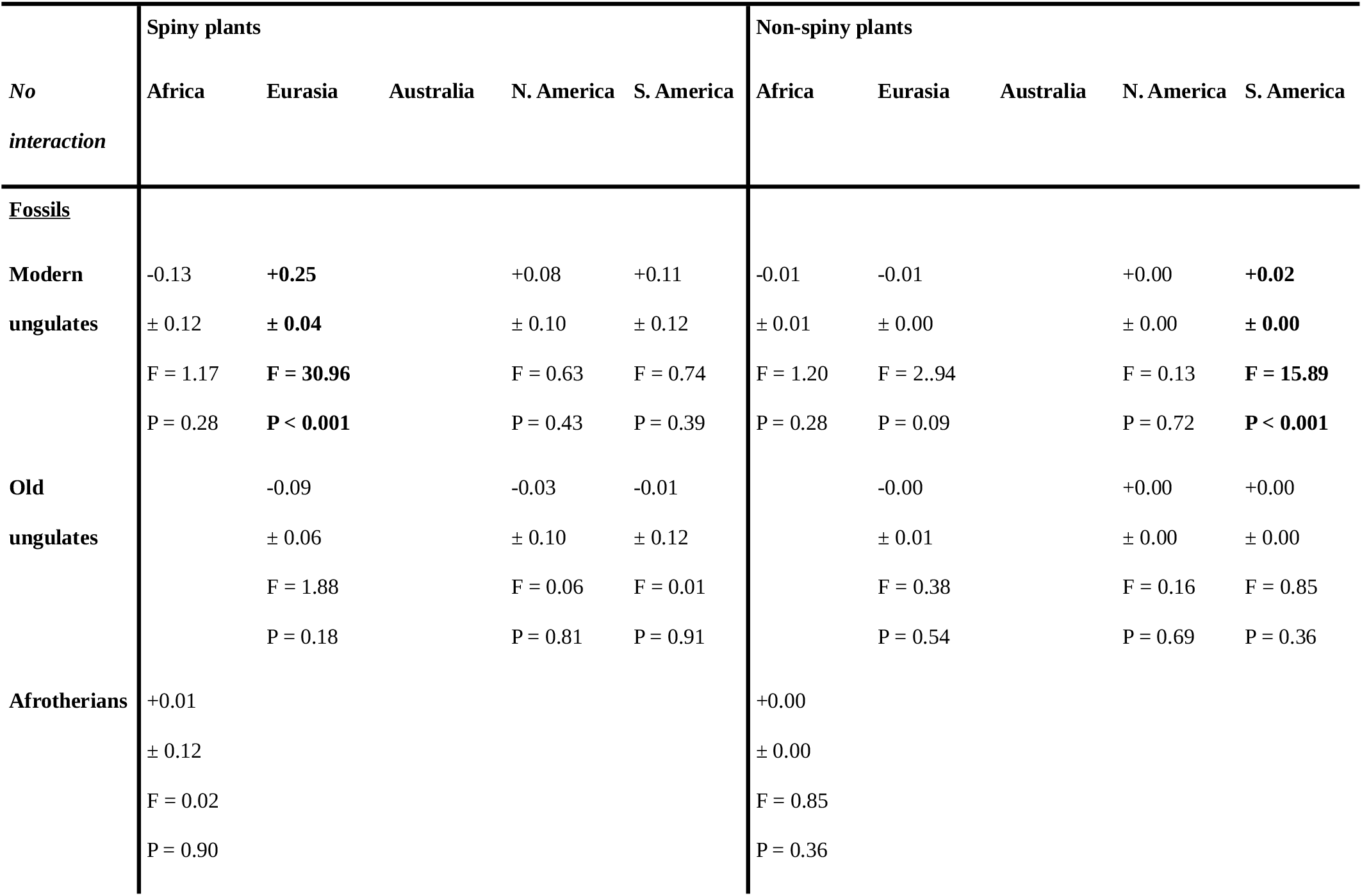
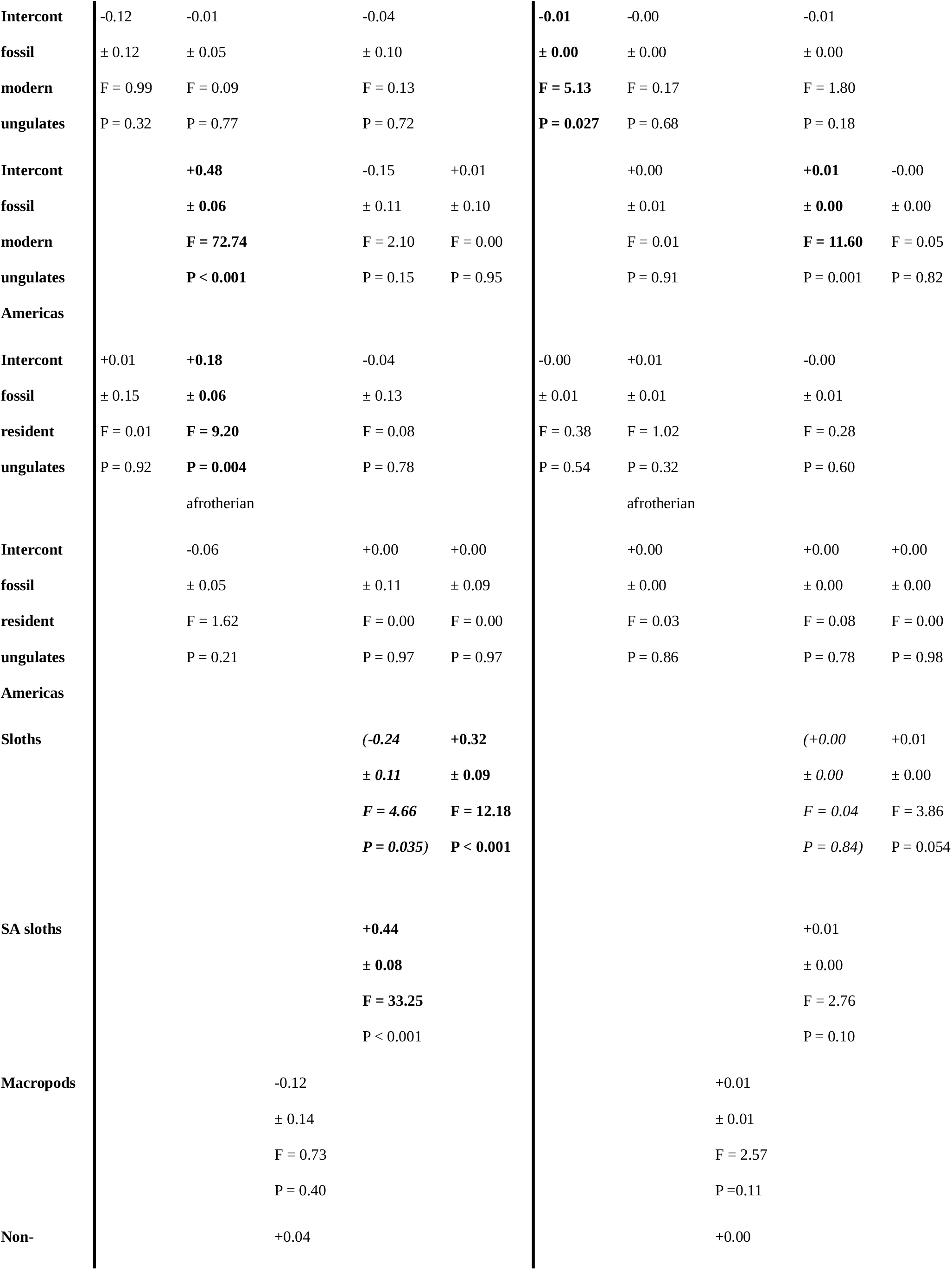
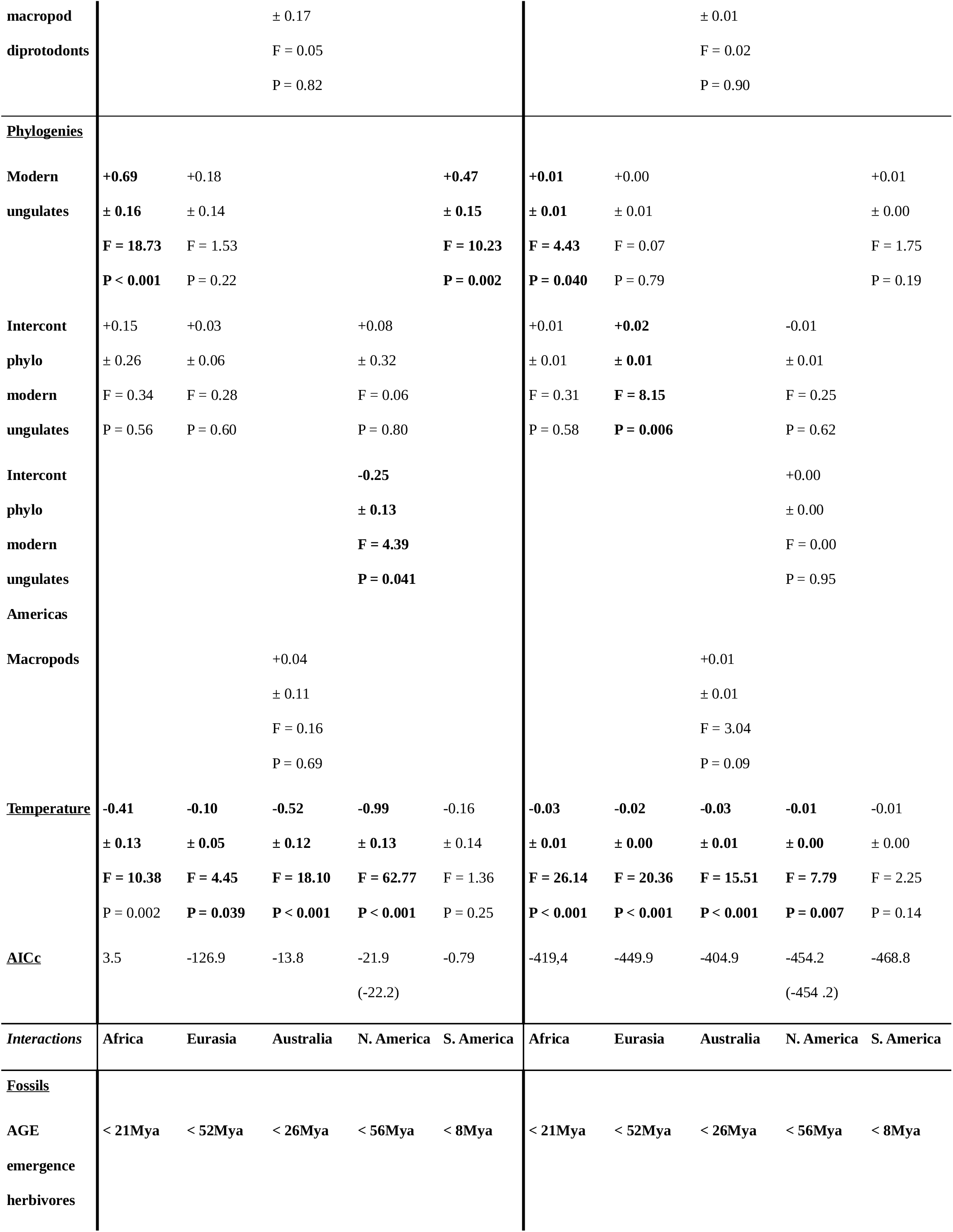
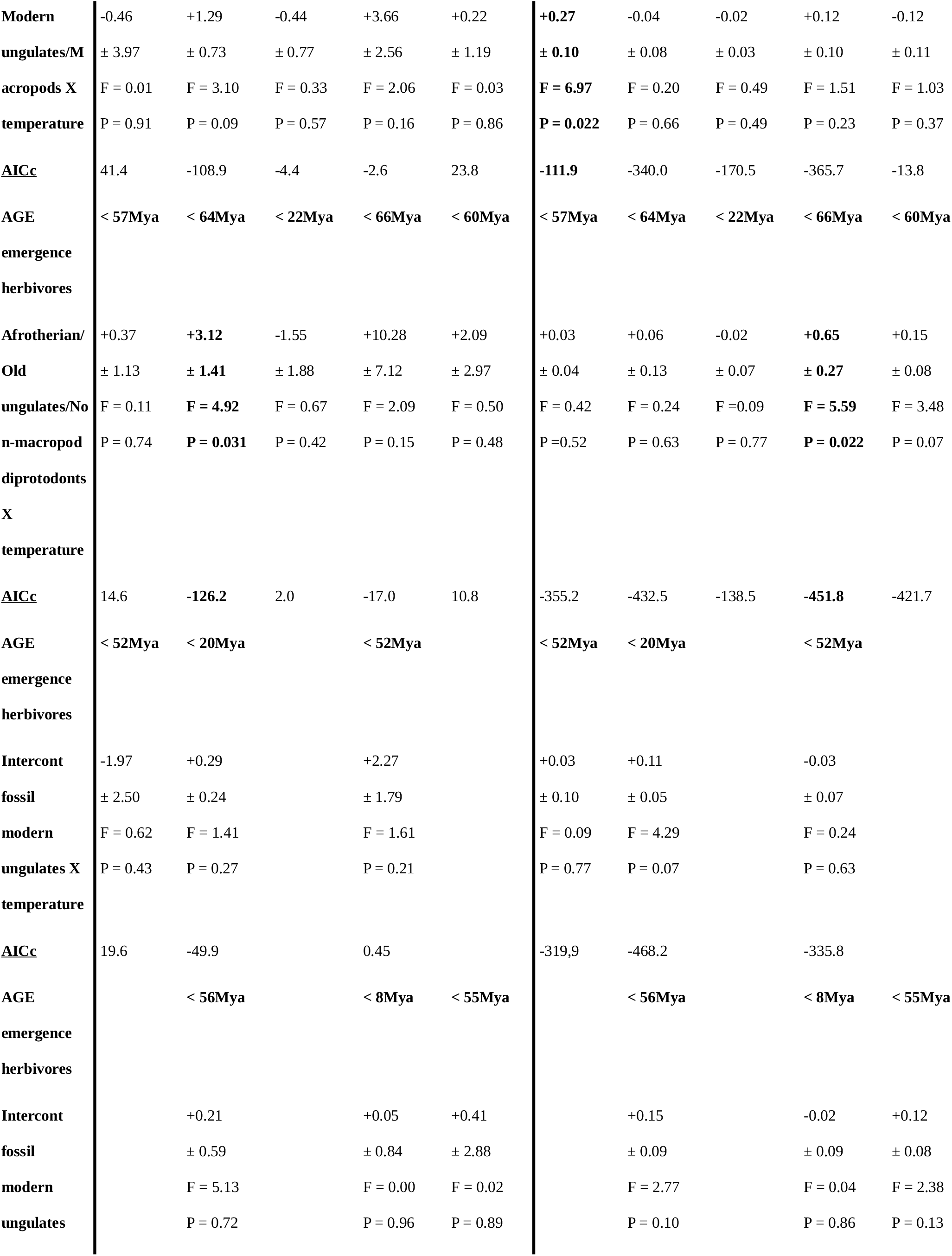
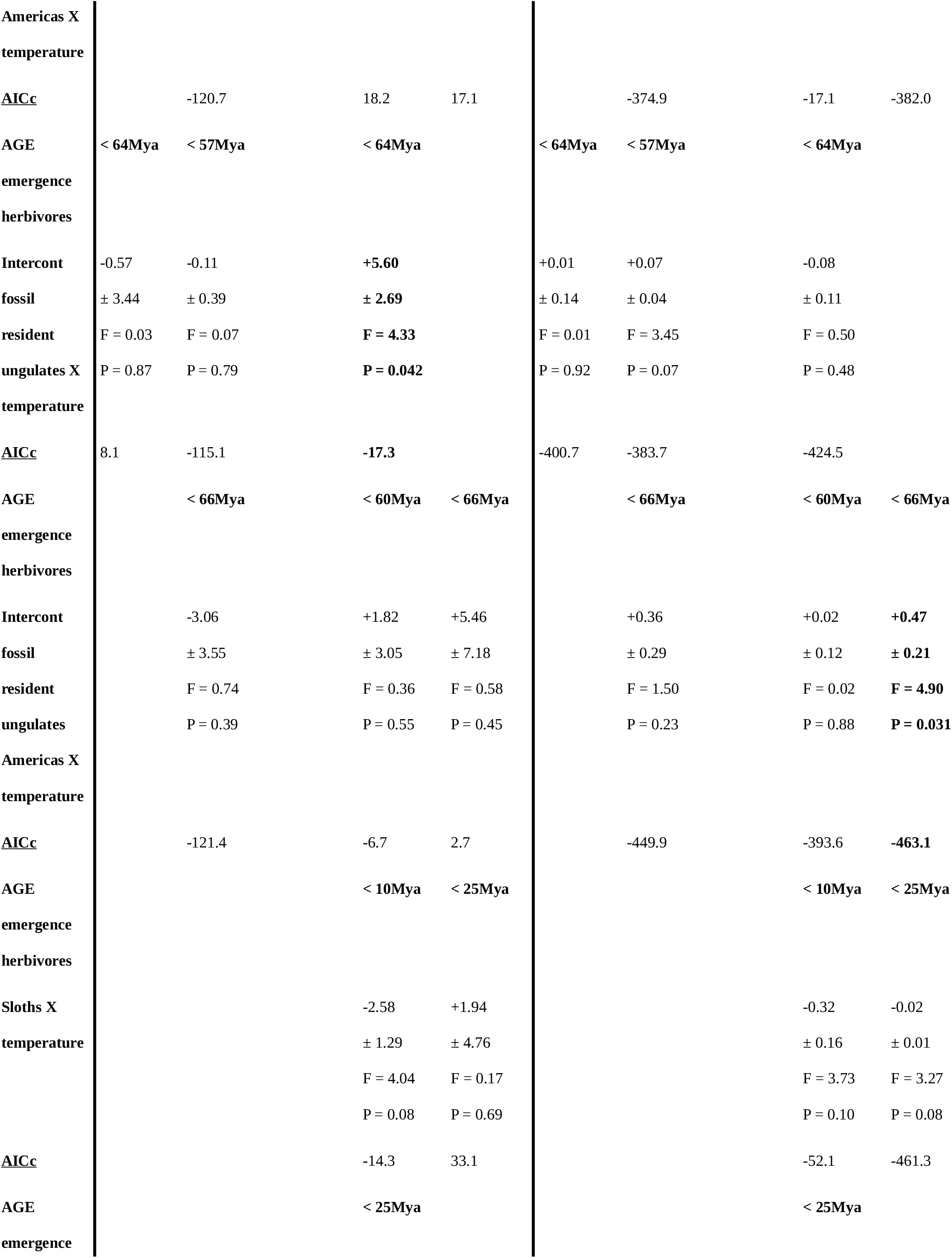
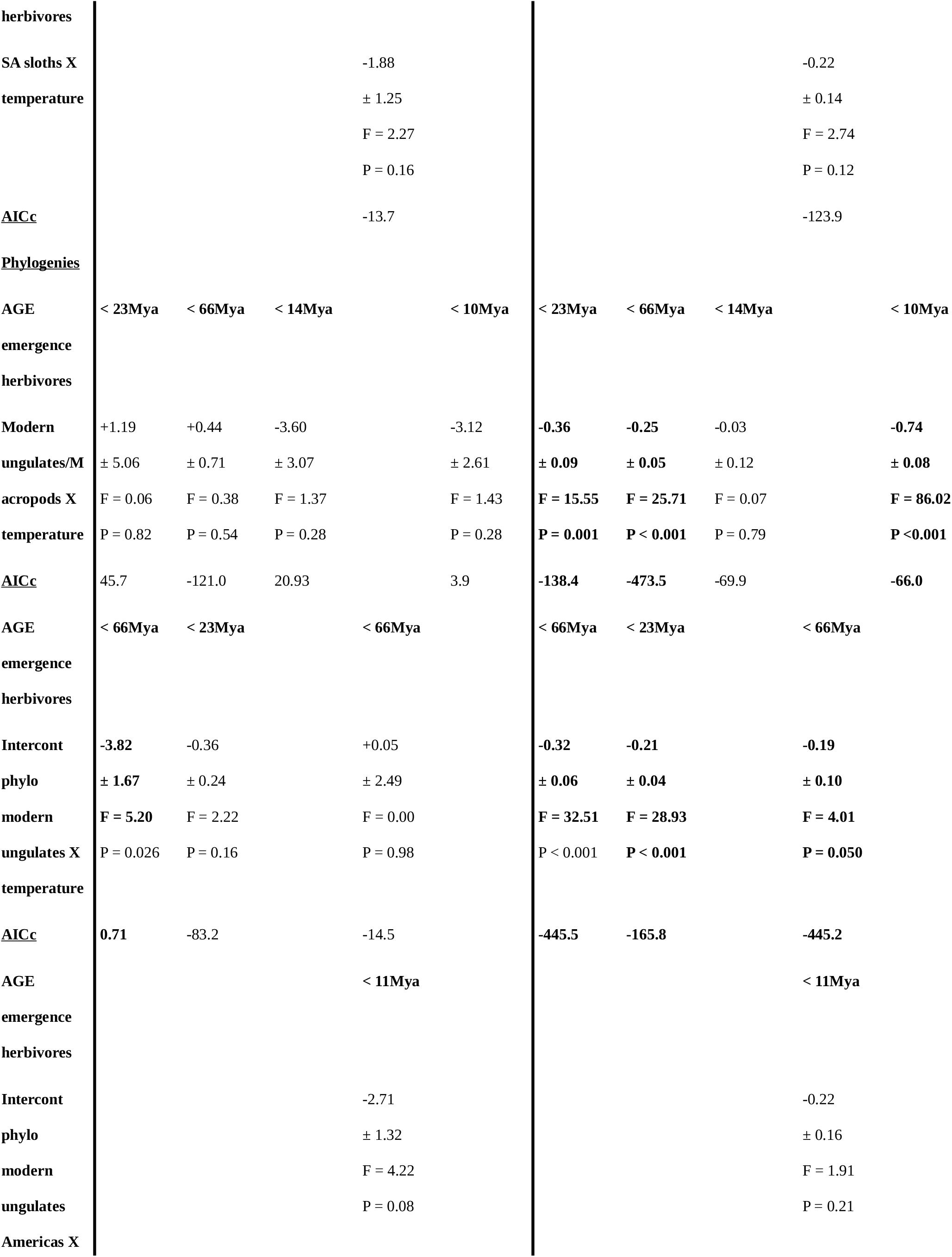
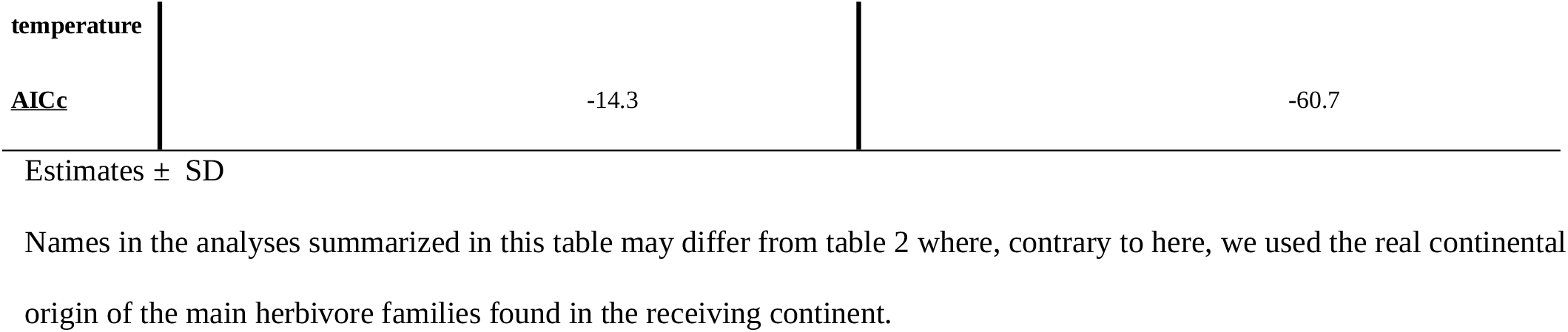
Estimates of linear models and AIC values testing the relationship between herbivore and plant (spiny and not spiny) yearly diversification rate, as well as global temperature for the Cenozoic in 5 continents, all single variables at a time (N = 67) and then one interaction at a time. Interaction was tested only starting from the first occurrence of a given mammal group, N then is the number of Mya + 1. Diversification of herbivores generally fits better to spiny plants while temperature fit better to non-spiny plants but the opposite pattern was found in Australia and North America; “migrant” herbivores appearing the more closely linked to spiny plant diversification but South American sloths and African Afrotherian may have also been involved. However, low temperature may increase the effect of herbivory by its effect on sea level and also possibly on plant growth, then increasing the number of predators coming from other continents through emerged lands and their impact on slowly growing plants.

**Table S5:**
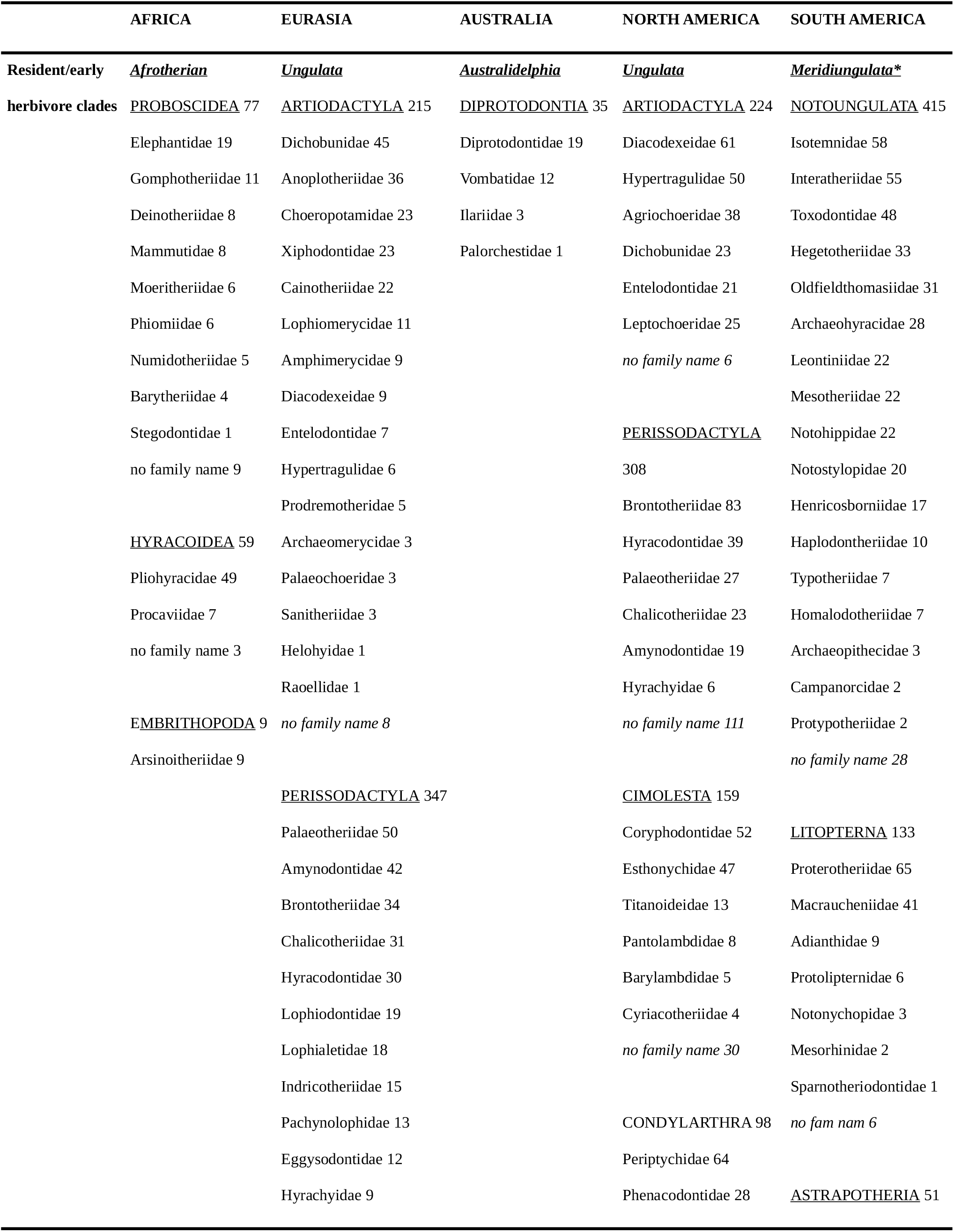
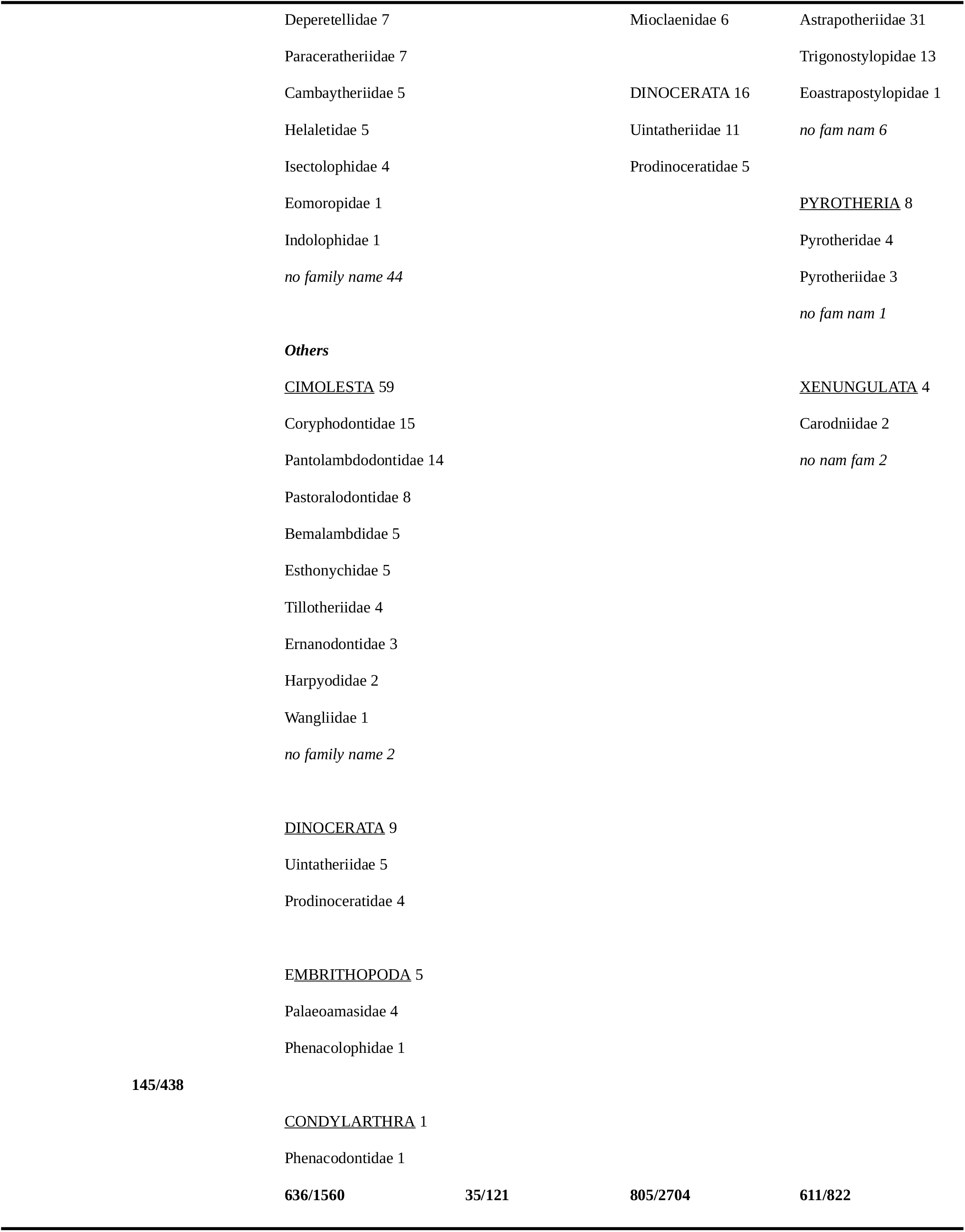
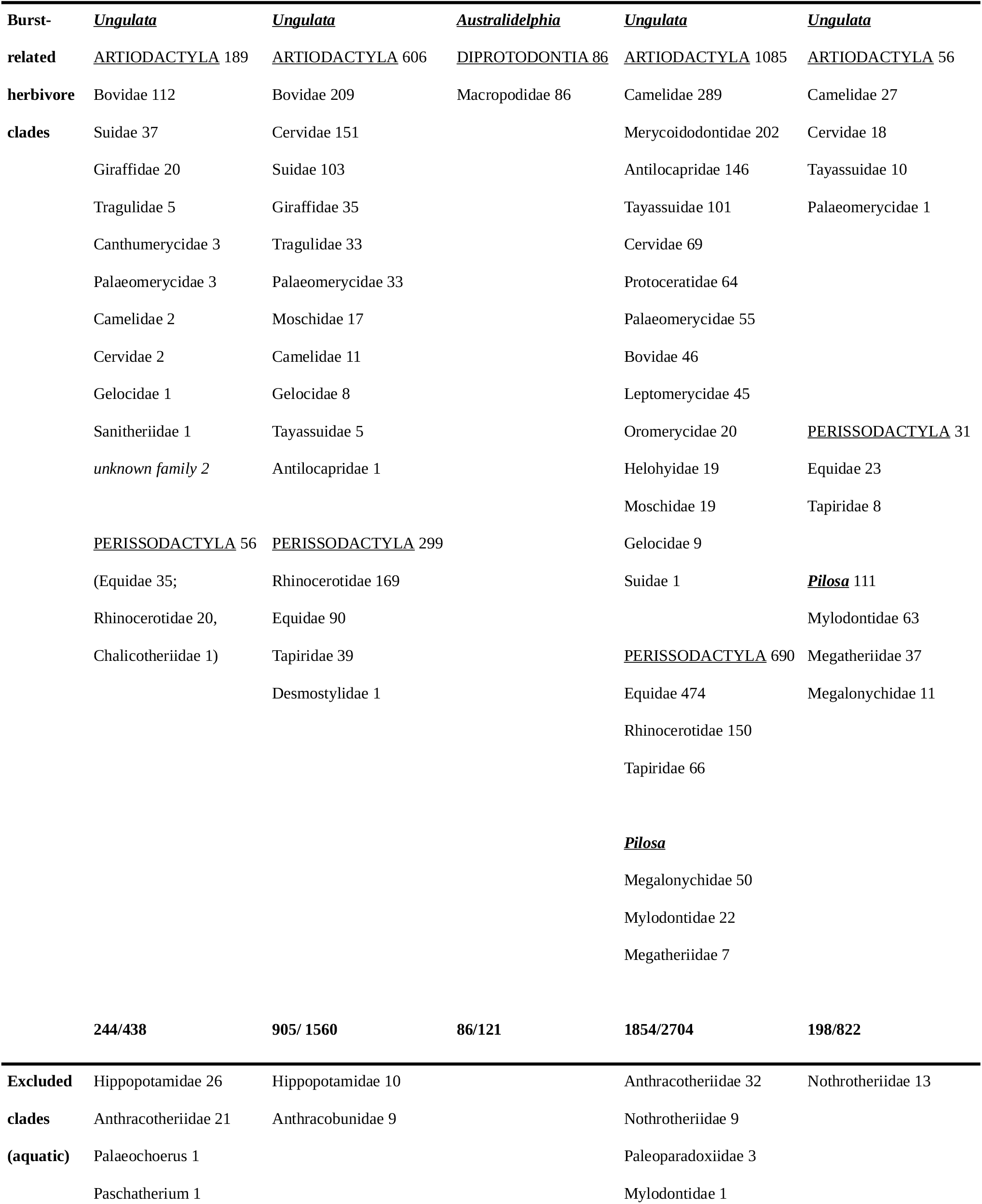
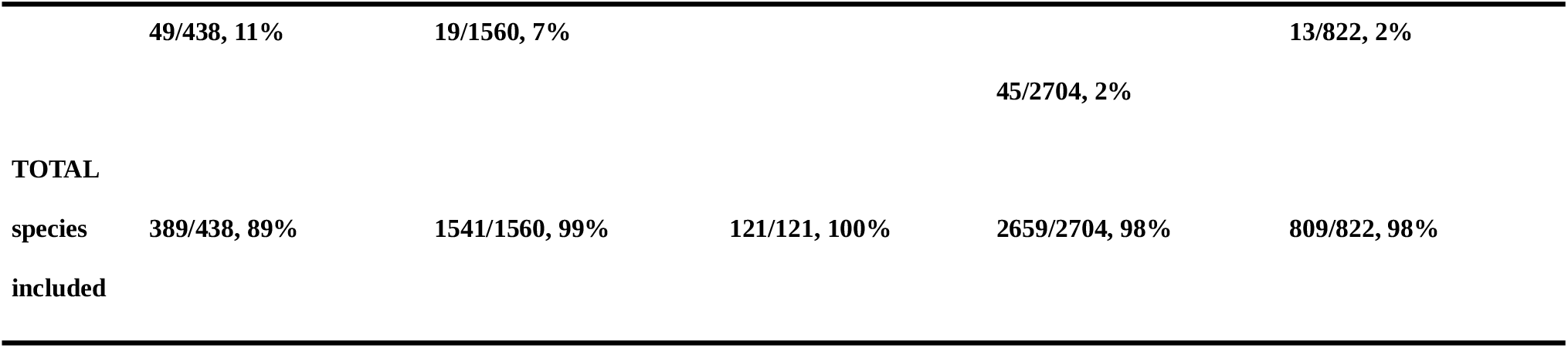
Details of mammalian herbivore clades included in our analyses

