## supplementary material for "The evolutionary history of spines – a Cenozoic arms race with mammals"

**This PDF file includes:**

Figs. S1 to S2

Tables S1 to S5

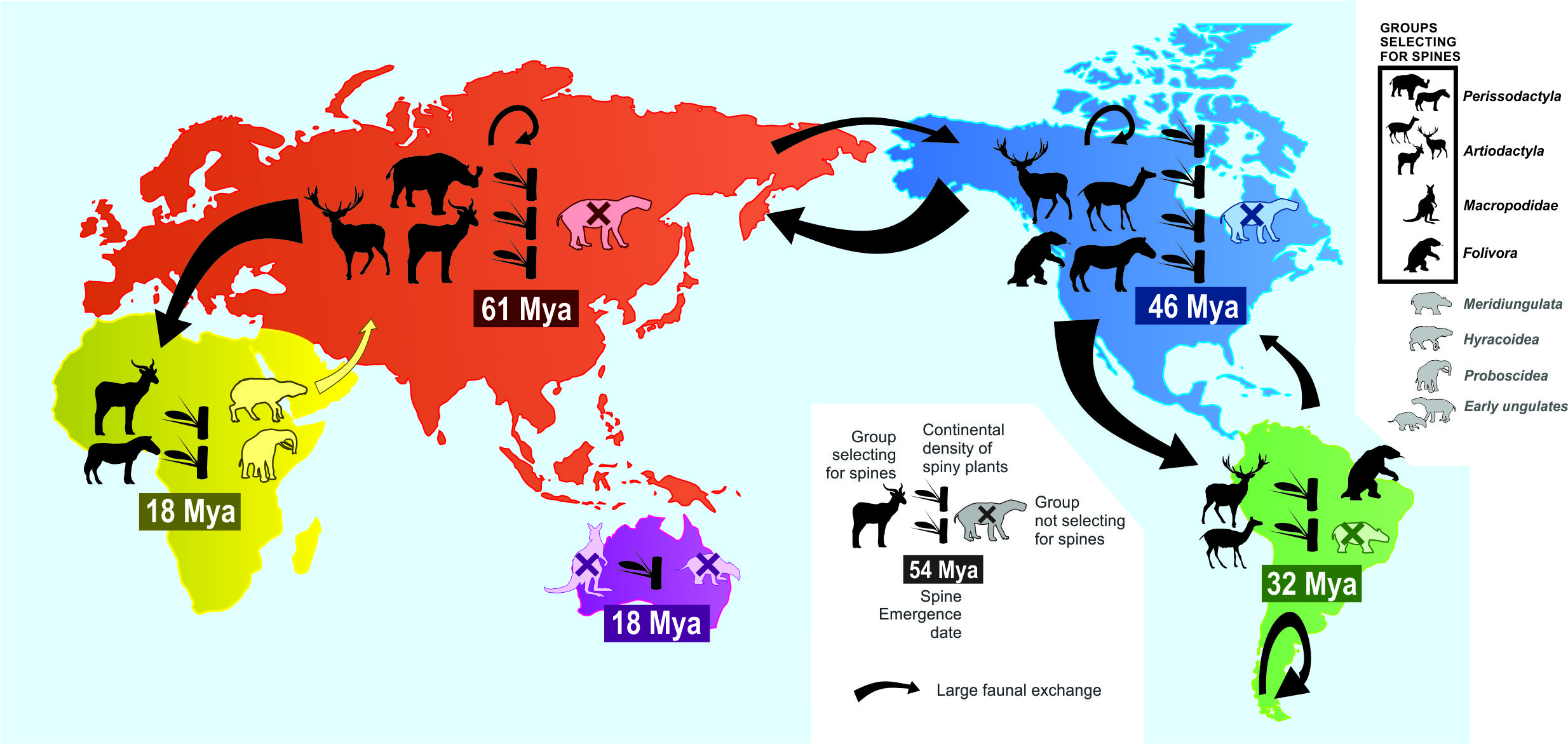
**Fig. S1:** Continental evolution and abundance of spiny plants has been mainly driven by speciation, migration and diversity of mammalian herbivores, especially Artiodactyla/Perissodactyla, and in North America, Folivora.

**Table S1:** Importance of correction for inflation of diversification rate (parameter Δ) in each continent. Corrected models including Δ always fitted the data better.

|  |  | **Africa** | **Eurasia** | **North**  **America** | **South**  **America** | **Australia** |
| --- | --- | --- | --- | --- | --- | --- |
| **Uncorrected model AIC** |  | **574** | **720** | **473** | **710** | **800** |
| **Corrected model AIC** |  | **513** | **688** | **402** | **639** | **703** |

**Table S2:** Relative fit of models fitting diversification rates to each continent (rows) using the correction for diversification inflation, Δ, either from the same continent or each of the other continents (columns); AIC values are given with AIC weights in parentheses. Emboldened values indicate best models in each row. Africa and North America show similar inflation of diversification rates, but other continents are best explained by their own Δ values.

|  | **Africa** (Δ=17) | **Eurasia** (Δ=9) | **Australia** (Δ=60) | **North America** (Δ=15) | **South America** (Δ=26) |
| --- | --- | --- | --- | --- | --- |
| **Africa** | **513** (0.45) | 520 (0.02) | 538 (0.00) | **514** (0.41) | 516 (0.13) |
| **Eurasia** | 699 (0.00) | **688** (0.97) | 790 (0.00) | 695 (0.03) | 718 (0.00) |
| **Australia** | 416 (0.00) | 430 (0.00) | **402** (0.98) | 418 (0.00) | 411 (0.02) |
| **North America** | **639** (0.43) | 644 (0.04) | 667 (0.00) | **639** (0.48) | 643 (0.06) |
| **South America** | 706 (0.18) | 722 (0.00) | 716 (0.00) | 708 (0.07) | **724** (0.75) |

**Table S3:** Estimates of linear models and AIC values testing the relationship between herbivore and plant (spiny and not spiny) yearly diversification rate and global temperature for the Cenozoic in 5 continents, one variable at a time (N = 67). Bold indicates the model fitting the best the data. Diversification of herbivores generally fit better to spiny plants while temperature fits better to non-spiny plants but the opposite pattern was found in Australia and North America; “migrant” herbivores appearing the more closely linked to spiny plant diversification but South American sloths and African Afrotherian may have also been involved.

|  | **Spiny plants** | | | | | **Non-spiny plants** | | | | |
| --- | --- | --- | --- | --- | --- | --- | --- | --- | --- | --- |
|  | **Africa** | **Eurasia** | **Australia** | **N. America** | **S. America** | **Africa** | **Eurasia** | **Australia** | **N. America** | **S. America** |
| **Fossils** |  |  |  |  |  |  |  |  |  |  |
| **Modern ungulates** | +0.50  ± 0.08  F = 39.39  P < 0.001 | +0.67  ± 0.07  F = 104.46  P < 0.001 |  | +0.34  ± 0.15  F = 5.06  P = 0.028 | +0.66  ± 0.10  F = 45.86  P < 0.001 | +0.01  ± 0.00  F = 13.79  P < 0.001 | +0.01  ± 0.00  F = 6.10  P = 0.016 |  | +0.00  ± 0.00  F = 0.62  P = 0.44 | **+0.03**  **± 0.00**  **F = 98.91**  **P < 0.001** |
| AIC | 30.8 | -37.6 |  | 64.3 | 24.7 | -393.0 | -404,3 |  | -405.5 | **-464.2** |
| **Old ungulates** |  | 0.21  ± 0.17  F = 1.41  P = 0.24 |  | -0.19  ± 0.15  F = 1.58  P = 0.21 | +0.10  ± 0.17  F = 0.37  P = 0.54 |  | -0.00  ± 0.01  F = 0.37  P = 0.54 |  | -0.00  ± 0.00  F = 0.17  P = 0.68 | +0.01  ± 0.01  F = 1.29  P = 0.26 |
| AIC |  | 25.2 |  | 67.7 | 60.1 |  | -398.6 |  | -405.1 | -403.6 |
| **Afrotherians** | +0.15  ± 0.15  F = 1.06  P = 0.31 |  |  |  |  | +0.00  ± 0.01  F = 0.81  P = 0.37 |  |  |  |  |
| AIC | 61.5 |  |  |  |  | -381.0 |  |  |  |  |
| **Intercont fossil modern ungulates** | +0.32  ± 0.13  F = 5.70  P = 0.020 | +0.18  ± 0.07  F = 6.11  P = 0.016 |  | +0.58  ± 0.13  F = 21.09  P < 0.001 |  | +0.01  ± 0.01  F = 2.22  P = 0.14 | +0.03  ± 0.01  F = 33.60  P < 0.001 |  | +0.01  ± 0.00  F = 2.22  P = 0.14 |  |
| AIC | 56.9 | 20.6 |  | 50.5 |  | -382.4 | -426.2 |  | -407.1 |  |
| **Intercont fossil modern ungulates Americas** |  | **+0.81**  **± 0.05**  **F = 233.62**  **P < 0.001** |  | +0.35  ± 0.10  F = 11.98  P < 0.001 | +0.16  ± 0.14  F = 1.28  P = 0.26 |  | +0.01  ± 0.00  F = 2.44  P = 0.12 |  | **+0.02**  **± 0.00**  **F = 78.46**  **P < 0.001** | +0.01  ± 0.00  F = 1.39  P = 0.24 |
| AIC |  | **-75.5** |  | 58.0 | 59.1 |  | -400.7 |  | **-457,9** | -403.7 |
| **Intercont fossil resident ungulates** | -0.06  ± 0.23  F = 0.07  P = 0.79 | +0.74  ± 0.07  F = 120.16  P < 0.001 |  | -0.10  ± 0.24  F = 0.18  P = 0.67 |  | +0.01  ± 0.01  F = 0.48  P = 0.49 | +0.01  ± 0.00  F = 2.76  P = 0.10 |  | +0.01  ± 0.01  F = 0.75  P = 0.39 |  |
| AIC | 62.5 | -43.5 |  | 69.2 |  | -380.6 | -401.0 |  | -405.7 |  |
| **Intercont fossil resident ungulates Americas** |  | -0.22  ± 0.11  F = 4.40  P = 0.040 |  | +0.23  ± 0.18  F = 1.62  P = 0.21 | -0.10  ± 0.14  F = 0.48  P = 0.49 |  | -0.00  ± 0.00  F = 0.18  P = 0.67 |  | +0.00  ± 0.01  F = 0.09  P = 0.77 | -0.00  ± 0.00  F = 0.57  P = 0.45 |
| AIC |  | 22.2 |  | 67.7 | 60.0 |  | -398.4 |  | -405 | -402.8 |
| **Sloths** |  |  |  | +0.51  ± 0.13  F = 14.12  P < 0.001 | +0.64  ± 0.09  F = 52.99  P < 0.001 |  |  |  | +0.02  ± 0.00  F = 46.05  P < 0.001 | +0.02  ± 0.00  F = 31.04  P < 0.001 |
| AIC |  |  |  | 56.2 | 20.5 |  |  |  | -440.8 | -428.4 |
| **SA sloths** |  |  |  | +0.75  ± 0.08  F = 78.57  P < 0.001 |  |  |  |  | +0.02  ± 0.00  F = 25.80  P < 0.001 |  |
| AIC |  |  |  | 16.3 |  |  |  |  | -427.3 |  |
| **Macropods** |  |  | +0.17  ± 0.07  F = 5.86  P = 0.018 |  |  |  |  | +0.03  ± 0.00  F = 57.08  P < 0.001 |  |  |
| AIC |  |  | 3.3 |  |  |  |  | -382.2 |  |  |
| **Non-macropod diprotodonts** |  |  | +0.23  ± 0.08  F = 7.69  P = 0.007 |  |  |  |  | +0.03  ± 0.00  F = 54.22  P < 0.001 |  |  |
| AIC |  |  | 1.6 |  |  |  |  | -380.7 |  |  |
| **Phylogenies** |  |  |  |  |  |  |  |  |  |  |
| **Modern ungulates** | **+0.73**  **± 0.08**  **F = 95.05**  **P < 0.001** | +0.13  ± 0.31  F = 0.18  P = 0.67 |  |  | **+0.86**  **± 0.10**  F = 81.24  **P < 0.001** | +0.02  ± 0.00  F = 30.73  P < 0.001 | +0.02  ± 0.01  F = 2.36  P = 0.13 |  |  | +0.03  ± 0.00  F = 72.41  P < 0.001 |
| AIC | **2,2** | 26,4 |  |  | **6.1** | -406.1 | -400.6 |  |  | -452.4 |
| **Intercont phylo modern ungulates** | +0.68  ± 0.40  F = 2.85  P = 0.10 | +0.25  ± 0.08  F = 9.00  P = 0.004 |  | +0.45  ± 0.43  F = 1.12  P = 0.29 |  | +0.02  ± 0.01  F = 1.86  P = 0.18 | +0.02  ± 0.00  F = 67.96  P < 0.001 |  | +0.01  ± 0.01  F = 1.45  P = 0.23 |  |
| AIC | 59,7 | 17,9 |  | 68.2 |  | -382.0 | -446.2 |  | -406.4 |  |
| **Intercont phylo modern ungulates Americas** |  |  |  | +0.53  ± 0.14  F = 14.49  P < 0.001 |  |  |  |  | +0.02  ± 0.00  F = 48.49  P < 0.001 |  |
| AIC |  |  |  | 55.9 |  |  |  |  | -442,2 |  |
| **Macropods** |  |  | +0.29  ± 0.08  F = 13.40  P < 0.001 |  |  |  |  | +0.04  ± 0.00  F = 58.26  P < 0.001 |  |  |
| AIC |  |  | -3,5 |  |  |  |  | -382.9 |  |  |
| **Temperature** | -0.82  ± 0.11  F = 53.05  P < 0.001 | -0.42  ± 0.10  F = 16.93  P < 0.001 | **-0.48**  **± 0.08**  **F = 34.57**  **P < 0.001** | -1.05  ± 0.09  F = 130.15  P < 0.001 | -0.80  ± 0.11  F = 50.95  P < 0.001 | **-0.03**  **± 0.00**  **F = 56.84**  **P < 0.001** | **-0.02**  **± 0.00**  **F = 70.04**  **P < 0.001** | **-0.05**  **± 0.00**  **F = 82.76**  **P < 0.001** | -0.03  ± 0.00  F = 61.06  P < 0.001 | -0.03  ± 0.00  F = 57.71  P < 0.001 |
| AIC | 22.6 | 11.1 | **-19.5** | -4.3 | 21.7 | **-422.2** | **-447.3** | **-395.0** | -449.3 | -444.8 |

Estimates ± SD

Names in the analyses summarized in this table may differ from table 2 where, contrary to here, we used the real continental origin of the main herbivore families found in the receiving continent.

**Table S4:** Estimates of linear models and AIC values testing the relationship between herbivore and plant (spiny and not spiny) yearly diversification rate, as well as global temperature for the Cenozoic in 5 continents, all single variables at a time (N = 67) and then one interaction at a time. Interaction was tested only starting from the first occurrence of a given mammal group, N then is the number of Mya + 1. Diversification of herbivores generally fits better to spiny plants while temperature fit better to non-spiny plants but the opposite pattern was found in Australia and North America; “migrant” herbivores appearing the more closely linked to spiny plant diversification but South American sloths and African Afrotherian may have also been involved. However, low temperature may increase the effect of herbivory by its effect on sea level and also possibly on plant growth, then increasing the number of predators coming from other continents through emerged lands and their impact on slowly growing plants.

|  | **Spiny plants** | | | | | **Non-spiny plants** | | | | |
| --- | --- | --- | --- | --- | --- | --- | --- | --- | --- | --- |
| ***No interaction*** | **Africa** | **Eurasia** | **Australia** | **N. America** | **S. America** | **Africa** | **Eurasia** | **Australia** | **N. America** | **S. America** |
| **Fossils** |  |  |  |  |  |  |  |  |  |  |
| **Modern ungulates** | -0.13  ± 0.12  F = 1.17  P = 0.28 | **+0.25**  **± 0.04**  **F = 30.96**  **P < 0.001** |  | +0.08  ± 0.10  F = 0.63  P = 0.43 | +0.11  ± 0.12  F = 0.74  P = 0.39 | -0.01  ± 0.01  F = 1.20  P = 0.28 | -0.01  ± 0.00  F = 2..94  P = 0.09 |  | +0.00  ± 0.00  F = 0.13  P = 0.72 | **+0.02**  **± 0.00**  **F = 15.89**  **P < 0.001** |
| **Old ungulates** |  | -0.09  ± 0.06  F = 1.88  P = 0.18 |  | -0.03  ± 0.10  F = 0.06  P = 0.81 | -0.01  ± 0.12  F = 0.01  P = 0.91 |  | -0.00  ± 0.01  F = 0.38  P = 0.54 |  | +0.00  ± 0.00  F = 0.16  P = 0.69 | +0.00  ± 0.00  F = 0.85  P = 0.36 |
| **Afrotherians** | +0.01  ± 0.12  F = 0.02  P = 0.90 |  |  |  |  | +0.00  ± 0.00  F = 0.85  P = 0.36 |  |  |  |  |
| **Intercont fossil modern ungulates** | -0.12  ± 0.12  F = 0.99  P = 0.32 | -0.01  ± 0.05  F = 0.09  P = 0.77 |  | -0.04  ± 0.10  F = 0.13  P = 0.72 |  | **-0.01**  **± 0.00**  **F = 5.13**  **P = 0.027** | -0.00  ± 0.00  F = 0.17  P = 0.68 |  | -0.01  ± 0.00  F = 1.80  P = 0.18 |  |
| **Intercont fossil modern ungulates Americas** |  | **+0.48**  **± 0.06**  **F = 72.74**  **P < 0.001** |  | -0.15  ± 0.11  F = 2.10  P = 0.15 | +0.01  ± 0.10  F = 0.00  P = 0.95 |  | +0.00  ± 0.01  F = 0.01  P = 0.91 |  | **+0.01**  **± 0.00**  **F = 11.60**  P = 0.001 | -0.00  ± 0.00  F = 0.05  P = 0.82 |
| **Intercont fossil resident ungulates** | +0.01  ± 0.15  F = 0.01  P = 0.92 | **+0.18**  **± 0.06**  **F = 9.20**  **P = 0.004**  afrotherian |  | -0.04  ± 0.13  F = 0.08  P = 0.78 |  | -0.00  ± 0.01  F = 0.38  P = 0.54 | +0.01  ± 0.01  F = 1.02  P = 0.32  afrotherian |  | -0.00  ± 0.01  F = 0.28  P = 0.60 |  |
| **Intercont fossil resident ungulates Americas** |  | -0.06  ± 0.05  F = 1.62  P = 0.21 |  | +0.00  ± 0.11  F = 0.00  P = 0.97 | +0.00  ± 0.09  F = 0.00  P = 0.97 |  | +0.00  ± 0.00  F = 0.03  P = 0.86 |  | +0.00  ± 0.00  F = 0.08  P = 0.78 | +0.00  ± 0.00  F = 0.00  P = 0.98 |
| **Sloths** |  |  |  | *(****-0.24***  ***± 0.11***  ***F = 4.66***  ***P = 0.035****)* | **+0.32**  **± 0.09**  **F = 12.18**  **P < 0.001** |  |  |  | *(+0.00*  *± 0.00*  *F = 0.04*  *P = 0.84)* | +0.01  ± 0.00  F = 3.86  P = 0.054 |
| **SA sloths** |  |  |  | **+0.44**  **± 0.08**  **F = 33.25**  P < 0.001 |  |  |  |  | +0.01  ± 0.00  F = 2.76  P = 0.10 |  |
| **Macropods** |  |  | -0.12  ± 0.14  F = 0.73  P = 0.40 |  |  |  |  | +0.01  ± 0.01  F = 2.57  P =0.11 |  |  |
| **Non-macropod diprotodonts** |  |  | +0.04  ± 0.17  F = 0.05  P = 0.82 |  |  |  |  | +0.00  ± 0.01  F = 0.02  P = 0.90 |  |  |
| **Phylogenies** |  |  |  |  |  |  |  |  |  |  |
| **Modern ungulates** | **+0.69**  **± 0.16**  **F = 18.73**  **P < 0.001** | +0.18  ± 0.14  F = 1.53  P = 0.22 |  |  | **+0.47**  **± 0.15**  **F = 10.23**  **P = 0.002** | **+0.01**  **± 0.01**  **F = 4.43**  **P = 0.040** | +0.00  ± 0.01  F = 0.07  P = 0.79 |  |  | +0.01  ± 0.00  F = 1.75  P = 0.19 |
| **Intercont phylo modern ungulates** | +0.15  ± 0.26  F = 0.34  P = 0.56 | +0.03  ± 0.06  F = 0.28  P = 0.60 |  | +0.08  ± 0.32  F = 0.06  P = 0.80 |  | +0.01  ± 0.01  F = 0.31  P = 0.58 | **+0.02**  **± 0.01**  **F = 8.15**  **P = 0.006** |  | -0.01  ± 0.01  F = 0.25  P = 0.62 |  |
| **Intercont phylo modern ungulates Americas** |  |  |  | **-0.25**  **± 0.13**  **F = 4.39**  **P = 0.041** |  |  |  |  | +0.00  ± 0.00  F = 0.00  P = 0.95 |  |
| **Macropods** |  |  | +0.04  ± 0.11  F = 0.16  P = 0.69 |  |  |  |  | +0.01  ± 0.01  F = 3.04  P = 0.09 |  |  |
| **Temperature** | **-0.41**  **± 0.13**  **F = 10.38**  P = 0.002 | **-0.10**  **± 0.05**  **F = 4.45**  **P = 0.039** | **-0.52**  **± 0.12**  **F = 18.10**  **P < 0.001** | **-0.99**  **± 0.13**  **F = 62.77**  **P < 0.001** | -0.16  ± 0.14  F = 1.36  P = 0.25 | **-0.03**  **± 0.01**  **F = 26.14**  **P < 0.001** | **-0.02**  **± 0.00**  **F = 20.36**  **P < 0.001** | **-0.03**  **± 0.01**  **F = 15.51**  **P < 0.001** | **-0.01**  **± 0.00**  **F = 7.79**  **P = 0.007** | -0.01  ± 0.00  F = 2.25  P = 0.14 |
| **AICc** | 3.5 | -126.9 | -13.8 | -21.9  (-22.2) | -0.79 | -419,4 | -449.9 | -404.9 | -454.2  (-454 .2) | -468.8 |
| ***Interactions*** | **Africa** | **Eurasia** | **Australia** | **N. America** | **S. America** | **Africa** | **Eurasia** | **Australia** | **N. America** | **S. America** |
| **Fossils** |  |  |  |  |  |  |  |  |  |  |
| **AGE emergence herbivores** | **< 21Mya** | **< 52Mya** | **< 26Mya** | **< 56Mya** | **< 8Mya** | **< 21Mya** | **< 52Mya** | **< 26Mya** | **< 56Mya** | **< 8Mya** |
| **Modern ungulates/Macropods X temperature** | -0.46  ± 3.97  F = 0.01  P = 0.91 | +1.29  ± 0.73  F = 3.10  P = 0.09 | -0.44  ± 0.77  F = 0.33  P = 0.57 | +3.66  ± 2.56  F = 2.06  P = 0.16 | +0.22  ± 1.19  F = 0.03  P = 0.86 | **+0.27**  **± 0.10**  **F = 6.97**  **P = 0.022** | -0.04  ± 0.08  F = 0.20  P = 0.66 | -0.02  ± 0.03  F = 0.49  P = 0.49 | +0.12  ± 0.10  F = 1.51  P = 0.23 | -0.12  ± 0.11  F = 1.03  P = 0.37 |
| **AICc** | 41.4 | -108.9 | -4.4 | -2.6 | 23.8 | **-111.9** | -340.0 | -170.5 | -365.7 | -13.8 |
| **AGE emergence herbivores** | **< 57Mya** | **< 64Mya** | **< 22Mya** | **< 66Mya** | **< 60Mya** | **< 57Mya** | **< 64Mya** | **< 22Mya** | **< 66Mya** | **< 60Mya** |
| **Afrotherian/Old ungulates/Non-macropod diprotodonts X temperature** | +0.37  ± 1.13  F = 0.11  P = 0.74 | **+3.12**  **± 1.41**  **F = 4.92**  **P = 0.031** | -1.55  ± 1.88  F = 0.67  P = 0.42 | +10.28  ± 7.12  F = 2.09  P = 0.15 | +2.09  ± 2.97  F = 0.50  P = 0.48 | +0.03  ± 0.04  F = 0.42  P =0.52 | +0.06  ± 0.13  F = 0.24  P = 0.63 | -0.02  ± 0.07  F =0.09  P = 0.77 | **+0.65**  **± 0.27**  **F = 5.59**  **P = 0.022** | +0.15  ± 0.08  F = 3.48  P = 0.07 |
| **AICc** | 14.6 | **-126.2** | 2.0 | -17.0 | 10.8 | -355.2 | -432.5 | -138.5 | **-451.8** | -421.7 |
| **AGE emergence herbivores** | **< 52Mya** | **< 20Mya** |  | **< 52Mya** |  | **< 52Mya** | **< 20Mya** |  | **< 52Mya** |  |
| **Intercont fossil modern ungulates X temperature** | -1.97  ± 2.50  F = 0.62  P = 0.43 | +0.29  ± 0.24  F = 1.41  P = 0.27 |  | +2.27  ± 1.79  F = 1.61  P = 0.21 |  | +0.03  ± 0.10  F = 0.09  P = 0.77 | +0.11  ± 0.05  F = 4.29  P = 0.07 |  | -0.03  ± 0.07  F = 0.24  P = 0.63 |  |
| **AICc** | 19.6 | -49.9 |  | 0.45 |  | -319,9 | -468.2 |  | -335.8 |  |
| **AGE emergence herbivores** |  | **< 56Mya** |  | **< 8Mya** | **< 55Mya** |  | **< 56Mya** |  | **< 8Mya** | **< 55Mya** |
| **Intercont fossil modern ungulates Americas X temperature** |  | +0.21  ± 0.59  F = 5.13  P = 0.72 |  | +0.05  ± 0.84  F = 0.00  P = 0.96 | +0.41  ± 2.88  F = 0.02  P = 0.89 |  | +0.15  ± 0.09  F = 2.77  P = 0.10 |  | -0.02  ± 0.09  F = 0.04  P = 0.86 | +0.12  ± 0.08  F = 2.38  P = 0.13 |
| **AICc** |  | -120.7 |  | 18.2 | 17.1 |  | -374.9 |  | -17.1 | -382.0 |
| **AGE emergence herbivores** | **< 64Mya** | **< 57Mya** |  | **< 64Mya** |  | **< 64Mya** | **< 57Mya** |  | **< 64Mya** |  |
| **Intercont fossil resident ungulates X temperature** | -0.57  ± 3.44  F = 0.03  P = 0.87 | -0.11  ± 0.39  F = 0.07  P = 0.79 |  | **+5.60**  **± 2.69**  **F = 4.33**  **P = 0.042** |  | +0.01  ± 0.14  F = 0.01  P = 0.92 | +0.07  ± 0.04  F = 3.45  P = 0.07 |  | -0.08  ± 0.11  F = 0.50  P = 0.48 |  |
| **AICc** | 8.1 | -115.1 |  | **-17.3** |  | -400.7 | -383.7 |  | -424.5 |  |
| **AGE emergence herbivores** |  | **< 66Mya** |  | **< 60Mya** | **< 66Mya** |  | **< 66Mya** |  | **< 60Mya** | **< 66Mya** |
| **Intercont fossil resident ungulates Americas X temperature** |  | -3.06  ± 3.55  F = 0.74  P = 0.39 |  | +1.82  ± 3.05  F = 0.36  P = 0.55 | +5.46  ± 7.18  F = 0.58  P = 0.45 |  | +0.36  ± 0.29  F = 1.50  P = 0.23 |  | +0.02  ± 0.12  F = 0.02  P = 0.88 | **+0.47**  **± 0.21**  **F = 4.90**  **P = 0.031** |
| **AICc** |  | -121.4 |  | -6.7 | 2.7 |  | -449.9 |  | -393.6 | **-463.1** |
| **AGE emergence herbivores** |  |  |  | **< 10Mya** | **< 25Mya** |  |  |  | **< 10Mya** | **< 25Mya** |
| **‍Sloths X temperature** |  |  |  | -2.58  ± 1.29  F = 4.04  P = 0.08 | +1.94  ± 4.76  F = 0.17  P = 0.69 |  |  |  | -0.32  ± 0.16  F = 3.73  P = 0.10 | -0.02  ± 0.01  F = 3.27  P = 0.08 |
| **AICc** |  |  |  | **-**14.3 | 33.1 |  |  |  | -52.1 | -461.3 |
| **AGE emergence herbivores** |  |  |  | **< 25Mya** |  |  |  |  | **< 25Mya** |  |
| **SA sloths X temperature** |  |  |  | -1.88  ± 1.25  F = 2.27  P = 0.16 |  |  |  |  | -0.22  ± 0.14  F = 2.74  P = 0.12 |  |
| **AICc** |  |  |  | -13.7 |  |  |  |  | -123.9 |  |
| **Phylogenies** |  |  |  |  |  |  |  |  |  |  |
| **AGE emergence herbivores** | **< 23Mya** | **< 66Mya** | **< 14Mya** |  | **< 10Mya** | **< 23Mya** | **< 66Mya** | **< 14Mya** |  | **< 10Mya** |
| **Modern ungulates/Macropods X temperature** | +1.19  ± 5.06  F = 0.06  P = 0.82 | +0.44  ± 0.71  F = 0.38  P = 0.54 | -3.60  ± 3.07  F = 1.37  P = 0.28 |  | -3.12  ± 2.61  F = 1.43  P = 0.28 | **-0.36**  **± 0.09**  **F = 15.55**  **P = 0.001** | **-0.25**  **± 0.05**  **F = 25.71**  **P < 0.001** | -0.03  ± 0.12  F = 0.07  P = 0.79 |  | **-0.74**  **± 0.08**  **F = 86.02**  **P <0.001** |
| **AICc** | 45.7 | -121.0 | 20.93 |  | 3.9 | **-138.4** | **-473.5** | -69.9 |  | **-66.0** |
| **AGE emergence herbivores** | **< 66Mya** | **< 23Mya** |  | **< 66Mya** |  | **< 66Mya** | **< 23Mya** |  | **< 66Mya** |  |
| **Intercont phylo modern ungulates X temperature** | **-3.82**  **± 1.67**  **F = 5.20**  P = 0.026 | -0.36  ± 0.24  F = 2.22  P = 0.16 |  | +0.05  ± 2.49  F = 0.00  P = 0.98 |  | **-0.32**  **± 0.06**  **F = 32.51**  P < 0.001 | **-0.21**  **± 0.04**  **F = 28.93**  **P < 0.001** |  | **-0.19**  **± 0.10**  **F = 4.01**  **P = 0.050** |  |
| **AICc** | **0.71** | -83.2 |  | -14.5 |  | **-445.5** | **-165.8** |  | **-445.2** |  |
| **AGE emergence herbivores** |  |  |  | **< 11Mya** |  |  |  |  | **< 11Mya** |  |
| **Intercont phylo modern ungulates Americas X temperature** |  |  |  | -2.71  ± 1.32  F = 4.22  P = 0.08 |  |  |  |  | -0.22  ± 0.16  F = 1.91  P = 0.21 |  |
| **AICc** |  |  |  | -14.3 |  |  |  |  | -60.7 |  |

Estimates ± SD

Names in the analyses summarized in this table may differ from table 2 where, contrary to here, we used the real continental origin of the main herbivore families found in the receiving continent.

**Table S5:** Details of mammalian herbivore clades included in our analyses

|  | **AFRICA** | **EURASIA** | **AUSTRALIA** | **NORTH AMERICA** | **SOUTH AMERICA** |
| --- | --- | --- | --- | --- | --- |
| **Resident/early herbivore clades** | ***Afrotherian***  PROBOSCIDEA 77 Elephantidae 19 Gomphotheriidae 11  Deinotheriidae 8 Mammutidae 8 Moeritheriidae 6 Phiomiidae 6 Numidotheriidae 5 Barytheriidae 4 Stegodontidae 1  no family name 9  HYRACOIDEA 59 Pliohyracidae 49 Procaviidae 7  no family name 3  EMBRITHOPODA 9 Arsinoitheriidae 9  **145/438** | ***Ungulata***  ARTIODACTYLA 215  Dichobunidae 45  Anoplotheriidae 36  Choeropotamidae 23  Xiphodontidae 23  Cainotheriidae 22  Lophiomerycidae 11  Amphimerycidae 9  Diacodexeidae 9  Entelodontidae 7  Hypertragulidae 6  Prodremotheridae 5  Archaeomerycidae 3  Palaeochoeridae 3  Sanitheriidae 3  Helohyidae 1  Raoellidae 1  *no family name 8*  PERISSODACTYLA 347  Palaeotheriidae 50  Amynodontidae 42 Brontotheriidae 34  Chalicotheriidae 31  Hyracodontidae 30 Lophiodontidae 19  Lophialetidae 18 Indricotheriidae 15 Pachynolophidae 13 Eggysodontidae 12  Hyrachyidae 9  Deperetellidae 7  Paraceratheriidae 7  Cambaytheriidae 5  Helaletidae 5  Isectolophidae 4  Eomoropidae 1  Indolophidae 1  *no family name 44*  ***Others***  CIMOLESTA 59  Coryphodontidae 15 Pantolambdodontidae 14  Pastoralodontidae 8 Bemalambdidae 5 Esthonychidae 5  Tillotheriidae 4  Ernanodontidae 3  Harpyodidae 2 Wangliidae 1  *no family name 2*  DINOCERATA 9  Uintatheriidae 5 Prodinoceratidae 4  EMBRITHOPODA 5 Palaeoamasidae 4  Phenacolophidae 1  CONDYLARTHRA 1  Phenacodontidae 1  **636/1560** | ***Australidelphia***  DIPROTODONTIA 35  Diprotodontidae 19 Vombatidae 12  Ilariidae 3  Palorchestidae 1  **35/121** | ***Ungulata***  ARTIODACTYLA 224  Diacodexeidae 61  Hypertragulidae 50  Agriochoeridae 38  Dichobunidae 23  Entelodontidae 21  Leptochoeridae 25  *no family name 6*  PERISSODACTYLA 308  Brontotheriidae 83  Hyracodontidae 39  Palaeotheriidae 27  Chalicotheriidae 23  Amynodontidae 19  Hyrachyidae 6  *no family name 111*  CIMOLESTA 159  Coryphodontidae 52  Esthonychidae 47  Titanoideidae 13  Pantolambdidae 8  Barylambdidae 5  Cyriacotheriidae 4  *no family name 30*  CONDYLARTHRA 98  Periptychidae 64 Phenacodontidae 28  Mioclaenidae 6  DINOCERATA 16  Uintatheriidae 11  Prodinoceratidae 5  **805/2704** | ***Meridiungulata****  NOTOUNGULATA 415  Isotemnidae 58  Interatheriidae 55  Toxodontidae 48  Hegetotheriidae 33  Oldfieldthomasiidae 31  Archaeohyracidae 28  Leontiniidae 22  Mesotheriidae 22  Notohippidae 22  Notostylopidae 20  Henricosborniidae 17  Haplodontheriidae 10  Typotheriidae 7  Homalodotheriidae 7  Archaeopithecidae 3  Campanorcidae 2  Protypotheriidae 2  *no family name 28*  LITOPTERNA 133  Proterotheriidae 65  Macraucheniidae 41  Adianthidae 9  Protolipternidae 6  Notonychopidae 3  Mesorhinidae 2  Sparnotheriodontidae 1  *no fam nam 6*  ASTRAPOTHERIA 51  Astrapotheriidae 31  Trigonostylopidae 13  Eoastrapostylopidae 1  *no fam nam 6*  PYROTHERIA 8  Pyrotheridae 4  Pyrotheriidae 3  *no fam nam 1*  XENUNGULATA 4  Carodniidae 2  *no nam fam 2*  **611/822** |

**Table S5. *Continued***

| **Burst-related herbivore clades** | ***Ungulata***  ARTIODACTYLA 189 Bovidae 112  Suidae 37  Giraffidae 20  Tragulidae 5 Canthumerycidae 3 Palaeomerycidae 3 Camelidae 2  Cervidae 2  Gelocidae 1  Sanitheriidae 1  *unknown family 2*  PERISSODACTYLA 56 (Equidae 35; Rhinocerotidae 20, Chalicotheriidae 1)  **244/438** | ***Ungulata***  ARTIODACTYLA 606 Bovidae 209  Cervidae 151  Suidae 103  Giraffidae 35  Tragulidae 33  Palaeomerycidae 33  Moschidae 17  Camelidae 11  Gelocidae 8  Tayassuidae 5  Antilocapridae 1  PERISSODACTYLA 299  Rhinocerotidae 169  Equidae 90  Tapiridae 39  Desmostylidae 1  **905/ 1560** | ***Australidelphia***  DIPROTODONTIA 86  Macropodidae 86  **86/121** | ***Ungulata***  ARTIODACTYLA 1085  Camelidae 289  Merycoidodontidae 202  Antilocapridae 146  Tayassuidae 101  Cervidae 69  Protoceratidae 64  Palaeomerycidae 55  Bovidae 46  Leptomerycidae 45  Oromerycidae 20  Helohyidae 19  Moschidae 19  Gelocidae 9  Suidae 1  PERISSODACTYLA 690  Equidae 474  Rhinocerotidae 150  Tapiridae 66  ***Pilosa***  Megalonychidae 50 Mylodontidae 22 Megatheriidae 7  **1854/2704** | ***Ungulata***  ARTIODACTYLA 56  Camelidae 27  Cervidae 18  Tayassuidae 10  Palaeomerycidae 1  PERISSODACTYLA 31  Equidae 23  Tapiridae 8  ***Pilosa*** 111  Mylodontidae 63 Megatheriidae 37 Megalonychidae 11  **198/822** |
| --- | --- | --- | --- | --- | --- |
| **Excluded clades (aquatic)** | Hippopotamidae 26  Anthracotheriidae 21  Palaeochoerus 1  Paschatherium 1  **49/438, 11%** | Hippopotamidae 10  Anthracobunidae 9  **19/1560, 7%** |  | Anthracotheriidae 32  Nothrotheriidae 9  Paleoparadoxiidae 3  Mylodontidae 1  **45/2704, 2%** | Nothrotheriidae 13  **13/822, 2%** |
| **TOTAL species included** | **389/438, 89%** | **1541/1560, 99%** | **121/121, 100%** | **2659/2704, 98%** | **809/822, 98%** |
